# Dynamical stability and accuracy of moment closure approximations for bursty gene expression with nonlinear degradation

**DOI:** 10.64898/2026.09.30.755644

**Authors:** Mohan Zhou, Chen Jia

## Abstract

Moment-closure methods provide a widely used approach for approximating low-order statistics of stochastic biochemical reaction networks, but the resulting closed equations may exhibit physically meaningless dynamics. Here, we systematically investigate the stability of second-order moment-closure approximations for a bursty gene expression model with nonlinear protein degradation. We consider the Gaussian, log-normal, and negative-binomial closures. By analyzing the resulting dynamical systems, we show that the Gaussian closure fails the proposed stability criterion for all parameter values, whereas both the log-normal and negative-binomial closures possess a unique physically accessible fixed point and are stable if and only if *kB*(1 + 2*B*) ≥ 1, where *B* is the mean burst size and *k* is the dimensionless protein synthesis rate. Moreover, this condition is automatically satisfied when the steady-state mean protein copy number is at least one, suggesting that it is mild in biologically relevant regimes. Numerical comparisons over a broad parameter range further show that the negative-binomial closure is the most accurate, while the Gaussian closure is the least accurate. We further develop higher-order negative-binomial moment-closure schemes and demonstrate that increasing the closure order substantially improves predictive accuracy. These results highlight the importance of considering dynamical stability, in addition to numerical accuracy, when assessing moment-closure approximations.

## 1 Introduction

Stochasticity in gene expression can give rise to pronounced cell-to-cell variability even within genetically identical populations [1, 2]. Over the past few decades, substantial progress has been made in both the theoretical and experimental study of stochastic gene expression at the single-cell level. Within the continuous-time Markovian framework, sample paths of mRNA and protein numbers can be generated numerically using the well-known stochastic simulation algorithm [3], while the time evolution of their probability distributions is governed by the chemical master equation (CME). In real biological systems, however, genes rarely function in isolation; instead, they interact with one another to form complex regulatory networks [4, 5]. Such gene interactions typically introduce nonlinear reaction propensities into stochastic gene expression models, rendering their dynamics considerably more difficult to analyze.

For simple stochastic gene expression models, the probability distributions of gene product numbers can sometimes be derived in closed form. Typical exactly solvable models include gene expression with two-state promoter switching [6–9], multi-state promoter switching [10–13], and autoregulation [14–18]. However, as gene regulatory mechanisms become increasingly intricate, analytical solutions rapidly become intractable, and considerable effort has therefore been devoted to solving the CME using numerical or approximate approaches [19–24]. In practical applications, however, we are sometimes less interested in the full probability distributions of gene product numbers than in their low-order moments and related statistics, such as the mean, variance, and correlation coefficient. In such cases, moment-based methods provide an alternative approach that directly describes the evolution of these statistics [25, 26]. For systems with nonlinear reaction propensities, the evolution of low-order moments generally depends on higher-order moments, giving rise to an unclosed hierarchy of moment equations. Moment-closure approximations address this problem by expressing the higher-order moments in terms of the retained lower-order moments, thereby yielding a finite system of differential equations.

To date, a wide variety of moment-closure methods have been developed. Distribution-based closure methods exploit the properties of a prescribed probability distribution to express high-order moments in terms of lower-order moments. Common distribution-based methods include the Poisson [27, 28], Gaussian [25, 29], log-normal [30, 31], gamma [32], and negative-binomial [33] closures. Maximum-entropy closures reconstruct a probability distribution by maximizing entropy subject to a set of low-order moment constraints and then calculate higher-order moments from the reconstructed distribution [34]. Conditional closure methods separate low-copy-number variables (such as promoter states) from high-copy-number variables (such as gene product numbers) and then apply conventional closure methods to the high-copy-number variables conditional on the discrete states of low-copy-number variables [35, 36]. Other commonly used moment-closure strategies include the linear mapping approximation [21], high-order linear mapping approximation [23], and variational approaches [37].

The quantitative performance of moment-closure methods is commonly assessed by comparing their predictions of low-order moments, such as means and variances, with numerical solutions of the CME. In particular, a well-behaved moment-closure method should yield physically meaningful results — whenever the CME admits a stationary solution, the moment-closure equations should possess a unique globally attractive fixed point, and their trajectories should exhibit non-negative means and variances for all times and all physically admissible initial conditions [29]. However, previous studies have shown that moment-closure equations can be numerically unstable and may give rise to unphysical behaviors, such as trajectories that diverge to infinity or enter regions with negative means or variances. Moreover, they may fail to correctly reproduce complex dynamical behaviors, such as multistability and sustained oscillations, exhibited by the underlying stochastic models [28, 32]. To date, these phenomena have been investigated primarily through numerical simulations, whereas rigorous mathematical studies of the dynamical properties of moment-closure equations remain very limited. In particular, a systematic understanding of the conditions under which different moment-closure methods yield numerically stable and physically meaningful dynamics is still lacking.

In the present paper, we systematically investigate the stability (also referred to as validity in Ref. [29]) of three moment-closure methods for a stochastic gene expression model with bursty protein production and nonlinear protein degradation. Here, a moment-closure method is said to be stable if it yields physically meaningful results for all times and all physically admissible initial conditions. The three methods considered are the Gaussian, log-normal, and negative-binomial closures. Our results show that the Gaussian closure is unstable for all parameter sets. Its moment-closure equations may admit zero, one, or two fixed points, and whenever fixed points exist, at least one of them is unstable. Consequently, physically meaningless trajectories necessarily arise, either diverging to infinity or entering regions with negative means or variances. In contrast, the moment-closure equations associated with both the log-normal and negative-binomial closures always possess a unique physically meaningful fixed point. Moreover, both methods are stable when the mean protein copy number of the underlying stochastic model exceeds one. Finally, we compare the accuracy of the three moment-closure methods through numerical simulations. We find that the Gaussian closure is the least accurate among the three methods, whereas the negative-binomial closure achieves the highest accuracy. We further extend the negative-binomial closure to higher-order schemes and show that retaining more moments leads to a substantial improvement in predictive accuracy.

The remainder of this paper is organized as follows. Section 2 derives the moment equations for a bursty gene expression model with nonlinear degradation and provides a rigorous mathematical definition of the stability of moment-closure methods. In Sections 3-5, we analyze and compare the stability of the Gaussian, log-normal, and negative-binomial closures. Section 6 further compares the accuracy of the three moment-closure strategies through numerical simulations. We conclude in Section 7.

## 2 Model and moment equations

Here we consider stochastic gene expression dynamics in a living cell with bursty protein synthesis and nonlinear protein degradation [29]. Let *G* denote the gene of interest and let *P* denote the corresponding protein. The reaction scheme is given by

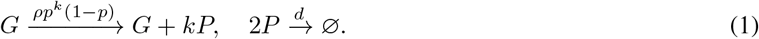

Here we assume that protein synthesis occurs in bursts with frequency *ρ* and random burst size *k* sampled from a geometric distribution with parameter *p* ∈ (0, 1), consistent with experimental observations [38]. The mean burst size is given by

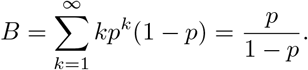

Moreover, we assume that proteins undergo post-translational dimerization followed by degradation. Experiments have shown that protein oligomerization can promote subsequent degradation and thus represents an important mechanism of post-translational regulation [39, 40]. Let *P*_*n*_(*t*) denote the probability of having *n* protein molecules in a single cell at time *t*, and let 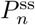 denote the corresponding steady-state probability. The degradation reaction is nonlinear because its propensity is proportional to *n*(*n ™* 1). The stochastic gene expression dynamics can then be described by a continuous-time Markov chain whose evolution is governed by the chemical master equation (CME)

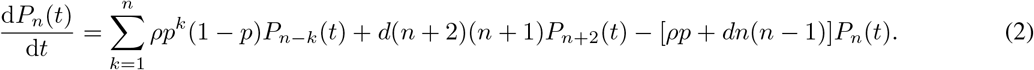

Using the standard Foster-Lyapunov ergodicity criterion for continuous-time Markov chains [41], it is straightforward to prove that the system is ergodic (see Appendix A for the proof). Specifically, regardless of the initial condition, the time-dependent distribution converges to the steady-state distribution as *t → ∞*, 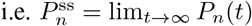.

We next focus on the first and second moments of protein numbers. Multiplying both sides of Eq. (2) by *n* and *n*^2^, respectively, and summing over *n* yields

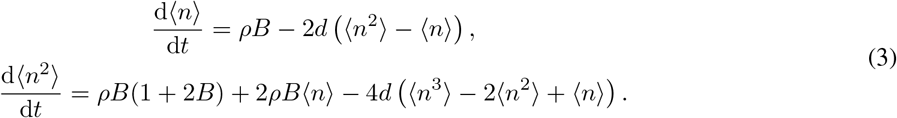

Note that this system of moment equations is not closed, i.e. the evolution of a moment of a given order depends on moments of higher orders, and hence an exact solution is generally unavailable. This difficulty arises from the nonlinear dependence of the propensity describing protein-protein interactions on protein numbers [30]. For convenience, we introduce the dimensionless time *τ* and dimensionless protein synthesis rate *k* as

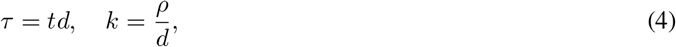

and further define

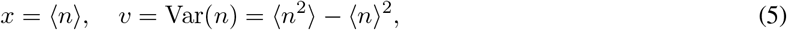

where *x* denotes the mean protein number and *v* denotes the variance of protein number fluctuations. In terms of the dimensionless time *τ*, Eq. (3) can be rewritten as the following system of differential equations:

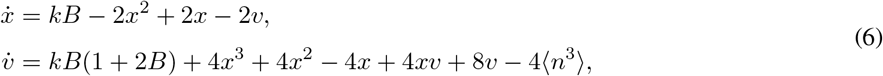

where 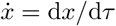 and 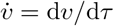 denote the derivatives of *x* and *v* with respect to *τ*.

Since the protein degradation reaction is nonlinear, Eq. (6) does not constitute a closed two-dimensional dynamical system for (*x, v*), because the second equation involves the third moment ⟨*n*^3^⟩ of protein numbers. A second-order moment-closure method provides an approximation of the form

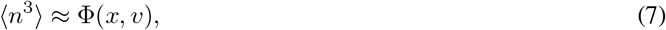

where Φ is referred to as the moment-closure function and is often determined by assuming that the protein number distribution belongs to a prescribed family, such as the Gaussian, log-normal, or other distributions. Substituting Eq. (7) into Eq. (6) yields the following closed planar dynamical system:

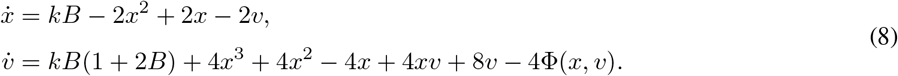

In the literature, this is often referred to as the *moment-closure approximation (MA) system* [29].

The performance of a moment-closure method is typically evaluated according to two complementary criteria: stability (also called validity in Ref. [29]) and accuracy. For a moment-closure method to be stable, the trajectories of the MA system should neither diverge to infinity nor produce negative means or variances (Fig. 1(a)). Previous studies have shown that the MA system associated with a moment-closure method may be unstable and exhibit three types of unphysical behavior (Fig. 1(b)): (i) the trajectories of the mean and variance may diverge to infinity (left panel); (ii) the trajectories may converge to an unphysical fixed point characterized by a negative mean or variance (middle panel); and (iii) although the trajectories eventually converge to a physically meaningful fixed point with positive mean and variance, the mean or variance may take negative values during the transient dynamics (right panel). To exclude these three types of unphysical behavior, we introduce the following definition.

**FIG. 1.**
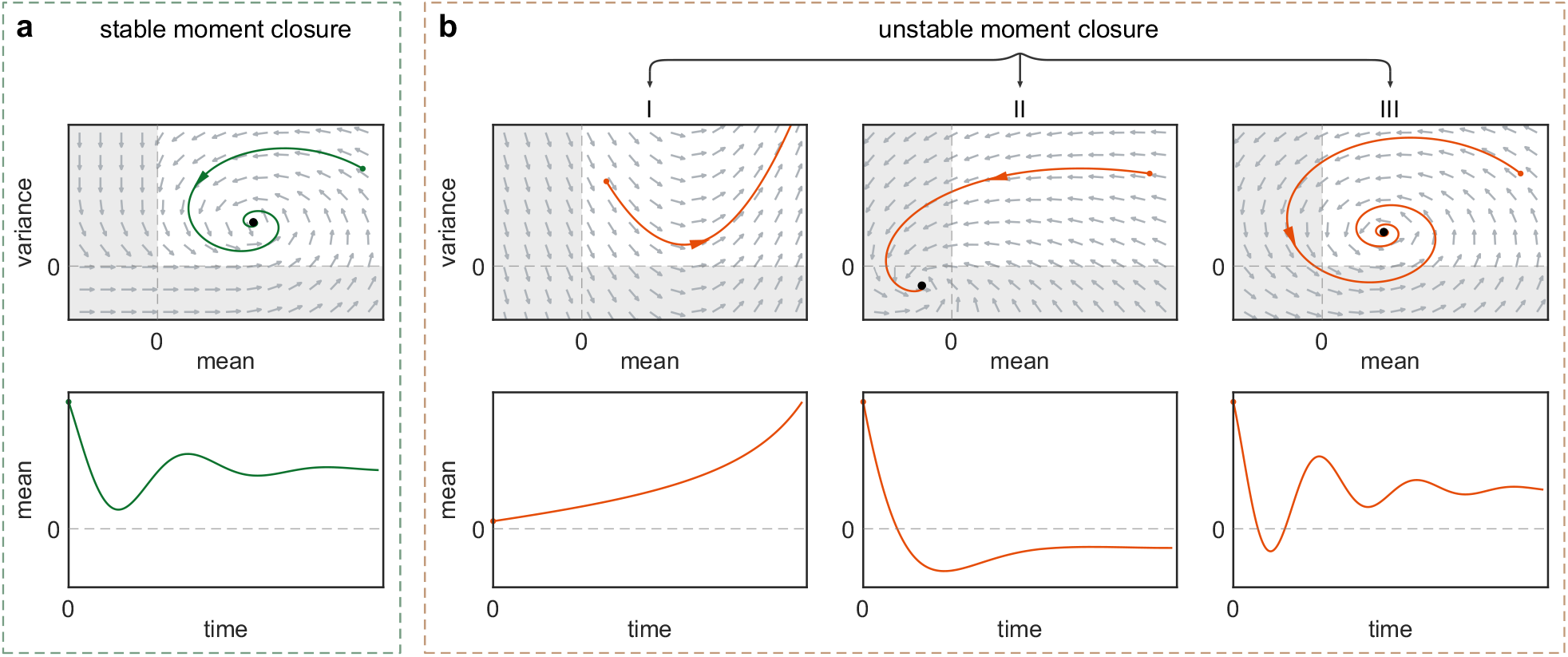
Schematic illustration of four typical dynamical behaviors of the MA system. **(a)** Typical phase portrait and trajectory of the mean for a stable moment-closure method. The white region in the (*x, v*)-phase portrait represents the physically meaningful region *D*_phys_, while the grey region represents the physically meaningless region. **(b)** Typical phase portraits and trajectories of the mean for an unstable moment-closure method. An unstable moment-closure method can exhibit three distinct types of unphysical behavior: (i) the trajectories of the mean and variance may diverge to infinity; (ii) the trajectories may converge to an unphysical fixed point characterized by a negative mean or variance; and (iii) although the trajectories eventually converge to a physically meaningful fixed point with positive mean and variance, the mean or variance may take negative values during the transient dynamics.

### Definition 1.

Consider the MA system given in Eq. (8) and let

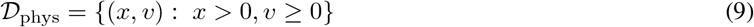

denote the physically meaningful region for the mean and variance. A fixed point (*x*_*∗*_, *v*_*∗*_) of the MA system is referred to as *physically accessible* or *physically meaningful* if (*x*_*∗*_, *v*_*∗*_) ∈ *D*_phys_. A moment-closure method is called *stable* if the following three conditions hold:

i. the MA system has a unique physically accessible fixed point (*x*_*∗*_, *v*_*∗*_) ∈ *D*_phys_;
ii. for any initial condition in *D*_phys_, the trajectory of the MA system does not blow up and remains within the physically meaningful region *D*_phys_, i.e. (*x*(*τ*), *v*(*τ*)) ∈ *D*_phys_ for all *τ* ≥ 0;
iii. for any initial condition in *D*_phys_, the trajectory of the MA system converges to the unique physically accessible fixed point, i.e. (*x*(*τ*), *v*(*τ*)) *→* (*x*_*∗*_, *v*_*∗*_) as *τ → ∞*.

Here, *D*_phys_ should be understood as the positivity region imposed by the basic physical requirements that both the mean and variance be nonnegative. Note that *D*_phys_ includes the boundary *v* = 0 but excludes *x* = 0. This is because the MA systems associated with many moment-closure methods, including the log-normal and negative-binomial closures considered below, become singular at *x* = 0. We therefore exclude *x* = 0 from *D*_phys_.

We emphasize that the stability of moment closures for the bursty gene expression model in Eq. (1) has previously been investigated in Ref. [29]. However, that study relied primarily on numerical investigations, whereas the present work provides a complete dynamical-systems analysis and derives precise mathematical conditions governing the stability of different moment-closure strategies. Moreover, the previous study was restricted to deterministic initial conditions (*v* = 0), while here we establish global dynamical properties of the MA systems under the more general setting of stochastic initial conditions (*v >* 0).

## 3 Stability for the Gaussian closure

We first investigate the Gaussian moment-closure method. Roughly speaking, the Gaussian closure approximates the protein number distribution by a Gaussian (normal) distribution. A key property of the Gaussian distribution is that all cumulants of order three and higher vanish [42]. Recall that the third-order cumulant of protein numbers is given by

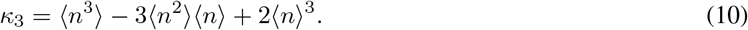

Under the Gaussian closure, we set *κ*_3_ = 0, and hence the third moment can be expressed in terms of the mean and variance as

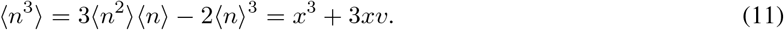

Substituting this expression into Eq. (6) yields the following MA system for the Gaussian closure:

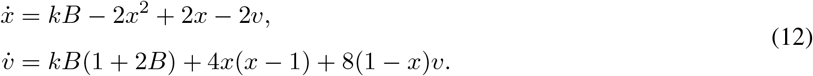

Recall that the MA system has two important parameters: the mean burst size *B* = *p/*(1 *™ p*) and the dimensionless protein synthesis rate *k* = *ρ/d*. The following proposition characterizes the number of physically accessible fixed points of the MA system and their stability in different parameter regimes of *B* and *k* (see Supplementary Sec. 1 for the proof).

### Proposition 1.

For any *B >* 0, the cubic equation

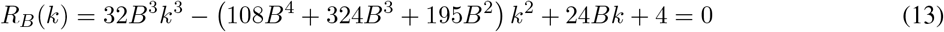

has two positive roots *k*_1_(*B*) *< k*_2_(*B*) and one negative root. For any *B >* 5*/*18, we have

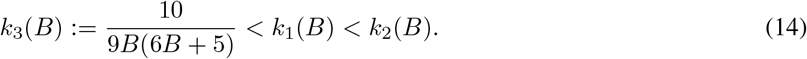

Moreover, the MA system given in Eq. (12) has the following properties:

i. When the dimensionless synthesis rate *k* ∈ (*k*_1_(*B*), *k*_2_(*B*)), the system possesses no physically accessible fixed points.
ii. When the dimensionless synthesis rate *k* = *k*_1_(*B*) or *k* = *k*_2_(*B*), the system possesses a unique physically accessible fixed point, which is unstable.
iii. When the mean burst size *B ≤* 5*/*18 and the dimensionless synthesis rate *k* ∈ (0, *k*_1_(*B*)) *∪* (*k*_2_(*B*), *∞*), the system possesses two physically accessible fixed points 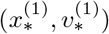 and 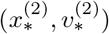 with 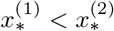. The fixed point 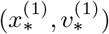 is a saddle, whereas 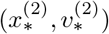 is locally asymptotically stable.
iv. When the mean burst size *B >* 5*/*18 and the dimensionless synthesis rate *k* ∈ (0, *k*_1_(*B*)) *∪* (*k*_2_(*B*), *∞*), the system possesses two physically accessible fixed points 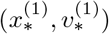 and 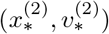 with 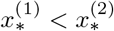. The fixed point 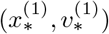 is a saddle, whereas 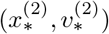 is locally asymptotically stable.
v. When the mean burst size *B >* 5*/*18 and the dimensionless synthesis rate *k* ∈ [*k*_3_(*B*), *k*_1_(*B*)), the system possesses two physically accessible fixed points 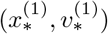 and 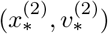 with 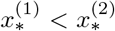. The fixed point 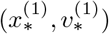 is a saddle, whereas 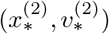 is an unstable node or focus.

According to the above proposition, except for the special cases *k* = *k*_1_(*B*) and *k* = *k*_2_(*B*), the MA system either has two physically accessible fixed points or has no physically accessible fixed point. When *k* = *k*_1_(*B*) or *k* = *k*_2_(*B*), the MA system possesses a unique physically accessible fixed point; however, this fixed point is unstable. All of these cases imply the instability of the Gaussian closure. We therefore obtain the following theorem.

### Theorem 1.

The Gaussian closure is unstable for all choices of *B* and *k*.

According to Proposition 1, the dynamical behavior of the Gaussian MA system can be classified into four distinct phases: (I) the system has no physically accessible fixed points; (II) the system has a unique physically accessible fixed point, which is unstable; (III) the system has two physically accessible fixed points, one stable and the other unstable; and (IV) the system has two physically accessible fixed points, both of which are unstable. To further characterize the parameter regions corresponding to these four phases, we illustrate the *B*-*k* phase diagram of the system in Fig. 2(a) and illustrate the typical phase portraits associated with the four phases in Fig. 2(b). Interestingly, when *B <* 5*/*18, only phases I-III can occur, separated by the two curves *k* = *k*_1_(*B*) and *k* = *k*_2_(*B*); when *B >* 5*/*18, all four phases can occur, separated by the three curves *k* = *k*_*i*_(*B*) for *i* = 1, 2, 3. This indicates that large burst sizes may give rise to richer dynamical behaviors. It is clear that phase I can only occur when the dimensionless protein synthesis rate *k* is neither too large nor too small, whereas phase III typically occurs when *k* is either very large or very small. Phase IV is confined to a very narrow region of the parameter space because the curves *k* = *k*_1_(*B*) and *k* = *k*_3_(*B*) are very close to each other. Among the four phases, only phase III can give rise to a large basin of attraction for a stable physically accessible fixed point (green region in Fig. 2(b)).

**FIG. 2.**
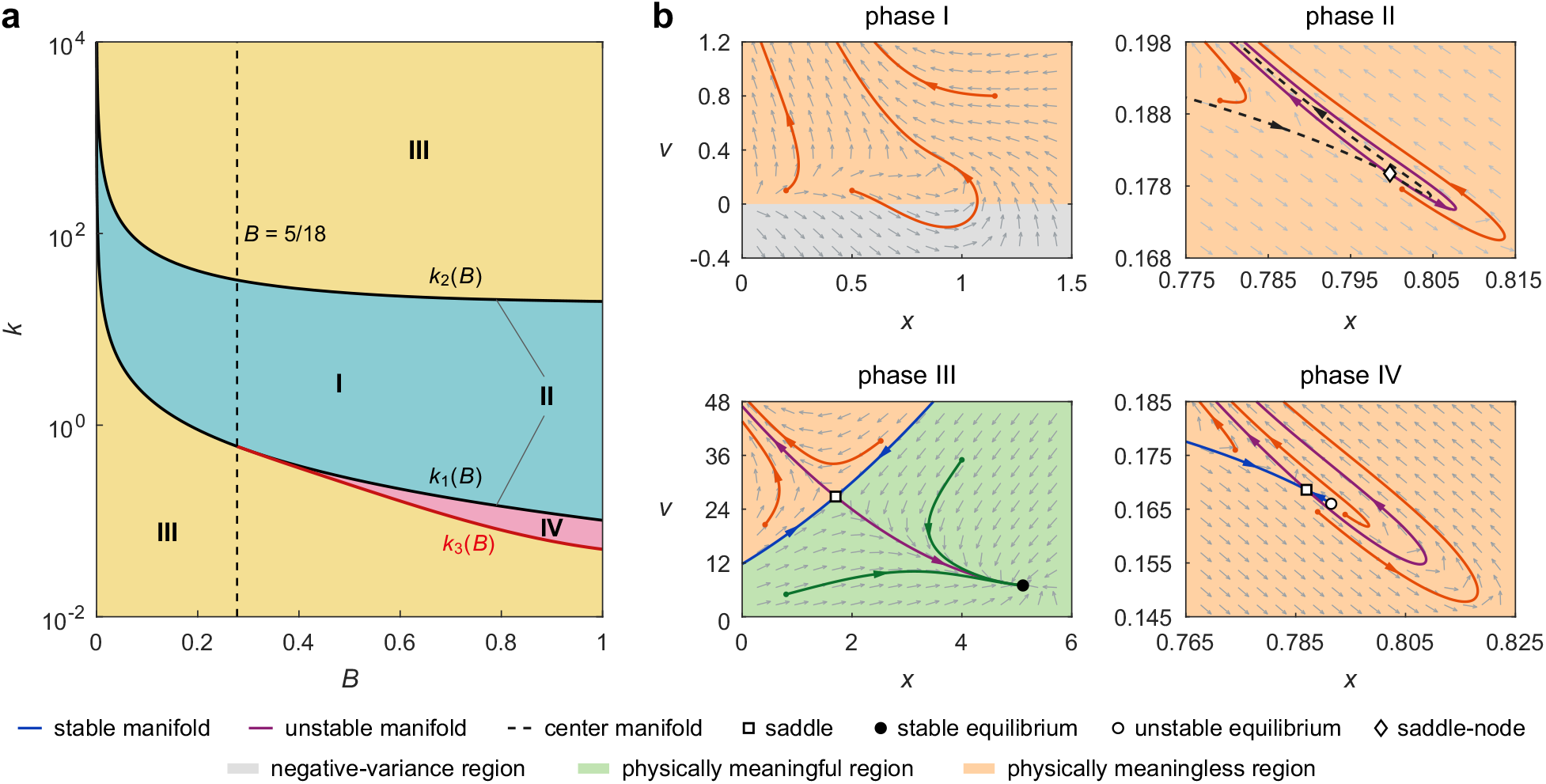
Phase diagram and typical phase portraits of the MA system for the Gaussian closure. **(a)** Phase diagram of the MA system in the *B*-*k* plane. The system can exhibit four distinct dynamical behaviors: (I) the system has no physically accessible fixed points; (II) the system has a unique physically accessible fixed point, which is unstable; (III) the system has two physically accessible fixed points, one stable and the other unstable; and (IV) the system has two physically accessible fixed points, both of which are unstable. When *B <* 5*/*18, only phases I-III can occur, separated by the two curves *k* = *k*_1_(*B*) and *k* = *k*_2_(*B*); when *B >* 5*/*18, all four phases can occur, separated by the three curves *k* = *k*_*i*_(*B*) for *i* = 1, 2, 3. **(b)** Typical phase portraits corresponding to the four phases. The model parameters are chosen as (*B, k*) = (0.4, 0.5) for phase I, (4, 0.009796) for phase II, (0.8, 70) for phase III, and (100, 1.9067 *×* 10^*™*5^) for phase IV, respectively.

**FIG. 3.**
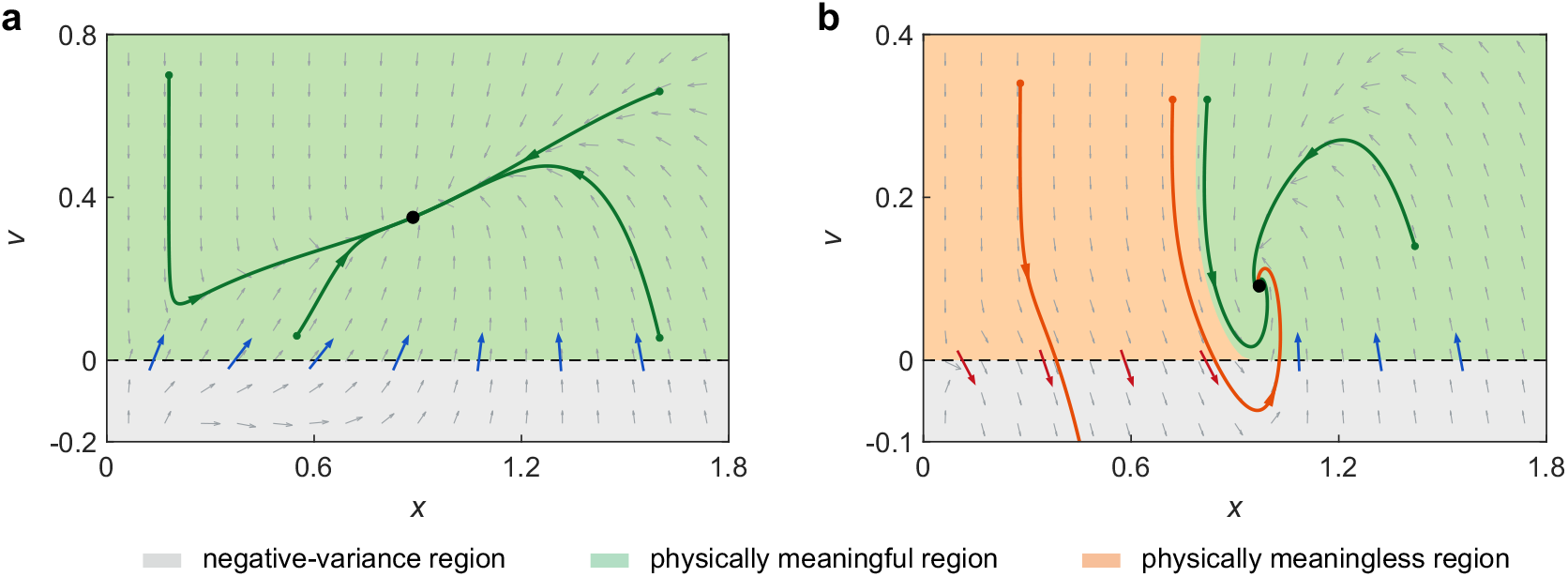
Typical phase portraits of the MA system for the log-normal closure. **(a)** The case *kB*(1 + 2*B*) *>* 1. The model parameters are chosen as *B* = 1.5, *k* = 1*/*3, and *kB*(1 + 2*B*) = 2. **(b)** The case *kB*(1 + 2*B*) *<* 1. The model parameters are chosen as *B* = 0.3, *k* = 5*/*12, and *kB*(1 + 2*B*) = 0.2. In (a) and (b), the grey region represents the negative-variance region. The green region indicates the set of initial conditions giving rise to physically meaningful trajectories (these trajectories remain within the physically meaningful region *D*_phys_ for all times and converge to the unique physically accessible fixed point). The orange region indicates the set of initial conditions giving rise to physically meaningless trajectories (these trajectories enter the negative-variance region at some time points).

## 4 Stability for the log-normal closure

We next focus on the log-normal moment-closure method, which is also referred to as the derivative-matching closure method [30, 31]. Roughly speaking, the log-normal closure approximates the protein number distribution by a log-normal distribution. Note that Gaussian random variables can take any real values, whereas log-normal random variables are restricted to positive values. Therefore, the log-normal closure may be more appropriate than the Gaussian closure for modeling protein numbers, which can only take nonnegative values.

By definition, a positive random variable *X* is said to be log-normally distributed if its logarithm *Z* = log(*X*) is normally distributed. Let *m*_*r*_ = E[*X*^*r*^] denote the *r*th-order moment of *X* and let *K*_*Z*_(*r*) = log E [*e*^*rZ*^] denote the cumulant generating function of *Z*. Since *X* = *e*^*Z*^, we have

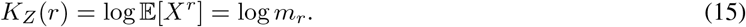

The cumulant generating function *K*_*Z*_(*r*) can be expanded as the power series

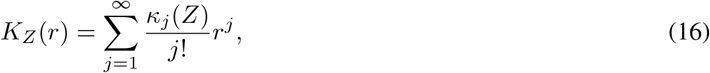

where *κ*_*j*_(*Z*) denotes the *j*th cumulant of *Z*. Since *Z* is Gaussian distributed, all its cumulants of order three and higher vanish, i.e. *κ*_*j*_(*Z*) = 0 for all *j* ≥ 3. Combining Eqs. (15) and (16), we can express log *m*_*r*_ in terms of the first two cumulants as

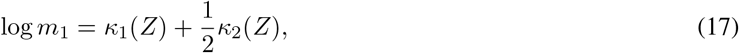

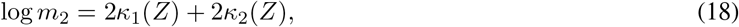

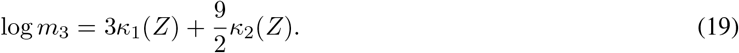

Combining Eqs. (17) and (18), we find that the first two cumulants can be expressed in terms of *m*_1_ and *m*_2_ as

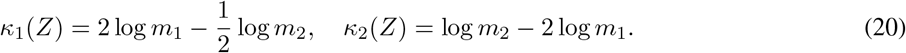

Substituting these expressions into Eq. (19) yields

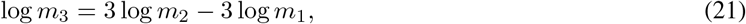

or, equivalently, 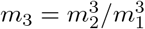. Thus, under the log-normal closure, the third moment can be approximated in terms of the mean and variance as

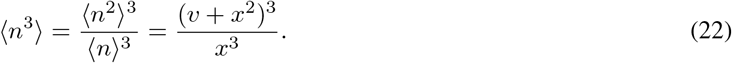

Substituting this expression into Eq. (6) yields the following MA system for the log-normal closure:

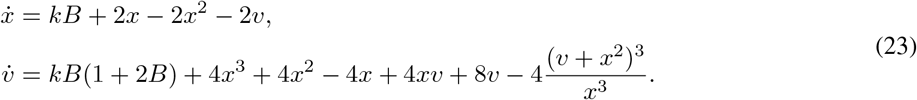

The following theorem characterizes the stability of the log-normal closure (see Supplementary Sec. 2 for the proof).

### Theorem 2.

For all choices of *B* and *k*, the log-normal MA system given in Eq. (23) possesses a unique physically accessible fixed point, which is locally asymptotically stable. Moreover, the log-normal closure is stable if and only if

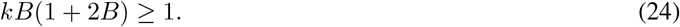

In other words, when *kB*(1 + 2*B*) ≥ 1, for any initial condition in *D*_phys_, the trajectory (*x*(*τ*), *v*(*τ*)) of the MA system remains in *D*_phys_ for all *τ* ≥ 0 and converges to the unique physically accessible fixed point.

We have shown that the Gaussian closure is unstable for the bursty gene expression model considered here, regardless of the choice of model parameters. In contrast, the log-normal closure is stable whenever *kB*(1 + 2*B*) ≥ 1. A natural question is whether this stability condition is typically satisfied in biologically relevant parameter regimes. This question is addressed by the following theorem.

### Theorem 3.

If the steady-state mean protein number *(n)*_ss_ ≥ 1 for the original stochastic model, then *kB*(1 + 2*B*) *>* 1.

*Proof*. Let *P* ^ss^(*n*) denote the steady-state probability of having *n* protein molecules, let *E* = {2*j* : *j* = 0, 1, 2, …} denote the set of all nonnegative even integers, and let *O* = {2*j* + 1 : *j* = 0, 1, 2, …} denote the set of all positive odd integers. Moreover, let

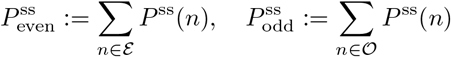

denote the steady-state probabilities that the protein copy number is even and odd, respectively. Let *q*_*nm*_ denote the transition rate of the Markovian model from microstate *n* to microstate *m*. It is clear that

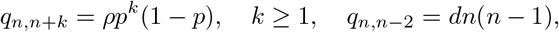

where the transition from microstate *n* to *n* + *k* is due to the synthesis reaction *G → G* + *kP*, whereas the transition from microstate *n* to *n ™* 2 is due to the degradation reaction 2*P →* ∅. Note that the degradation reaction does not change the parity of the protein number, whereas the synthesis reaction changes its parity only when the burst size *k* is odd. Hence, the total probability flow from the set ε to the set *O* is given by

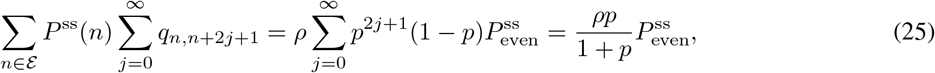

and the total probability flow from the set *O* to the set *ε* is given by

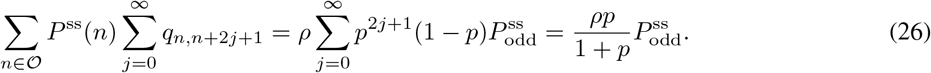

At steady state, the total probability flows between *E* and *O* must balance. Hence,

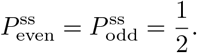

Moreover, for every nonnegative integer *n*, it is straightforward to verify that

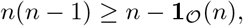

where **1**_*O*_ denotes the indicator function of the set *O*, which takes the value 1 when *n* ∈ *O* and 0 when *n* ∈ *ε*. It follows that

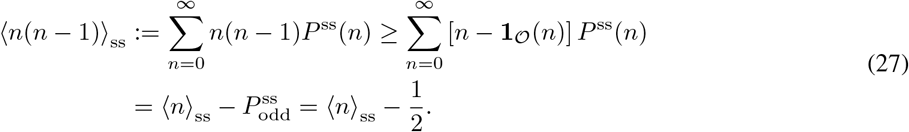

Setting the time derivative to zero in the first equation of Eq. (3), we obtain

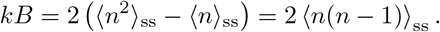

It thus follows from Eq. (27) that

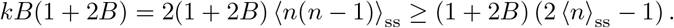

Therefore, *(n)*_ss_ ≥ 1 implies that

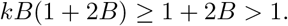

This completes the proof.

The above theorem links the condition *kB*(1 + 2*B*) ≥ 1 to the mean protein number of the original stochastic dynamics. Table 1 summarizes the median mRNA and protein copy numbers per cell across different cell types reported in the literature. While the median mRNA copy number is less than 1 per cell in bacteria, it is generally greater than 1 in yeast and higher eukaryotic cells. In contrast, the median protein copy number is substantially greater than 1 across all cell types, ranging from bacteria to yeast and higher eukaryotes. This suggests that *kB*(1 + 2*B*) ≥ 1 is a mild condition that is expected to be widely satisfied in biological systems.

**Table 1.** Representative median mRNA and protein abundances in different cell types. For *Mycoplasma pneumoniae*, the values reported in [45] are means rather than medians.

| cell type | median mRNA copies per cell | median protein copies per cell | Reference |
| --- | --- | --- | --- |
| *Escherichia coli* | 0.064 | 18.2 | [43, 44] |
| *Mycoplasma pneumoniae* | 0.04 | 167 | [45] |
| *Saccharomyces cerevisiae* (budding yeast) | 1.2 | 2,460 | [46] |
| *Schizosaccharomyces pombe* (fission yeast) | 2.4 | 3,919 | [47] |
| mouse fibroblasts (NIH3T3) | 17 | 50,000 | [48] |
| HeLa cells | NA | 18,000 | [49] |

## 5 Stability for the negative-binomial closure

We finally examine the negative-binomial moment-closure method [33]. Numerous single-cell experiments have shown that mRNA and protein number distributions in living cells can be well described by negative-binomial distributions [43, 50]. Note that log-normal random variables can take any positive real values but cannot take the value zero, whereas negative-binomial random variables are restricted to nonnegative integer values. Therefore, the negative-binomial closure may be more appropriate than the log-normal closure for modeling protein numbers since (i) protein numbers are always integer-valued and (ii) they can take the value zero with positive probability.

The basic idea of the negative-binomial closure is to approximate the protein number distribution by a negative-binomial distribution. Recall that a nonnegative integer-valued random variable *X* follows a negative-binomial distribution with parameters *α, β >* 0 if

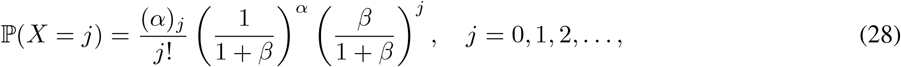

where (*α*)_*j*_ = *α*(*α* + 1) *…* (*α* + *j ™* 1) denotes the Pochhammer symbol. Moreover, let

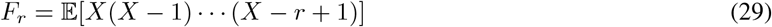

denote the *r*th-order factorial moment of *X*, with the convention *F*_0_ = 1. If *X* is negative-binomially distributed, it is straightforward to verify that *F*_*r*_ = *β*^*r*^(*α*)_*r*_ [51]. It follows immediately that

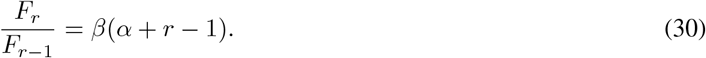

In Appendix B, we prove the following two combinatorial identities: for any integer *q* ≥ 3, we have

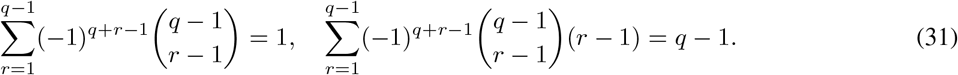

Combining these two identities yields

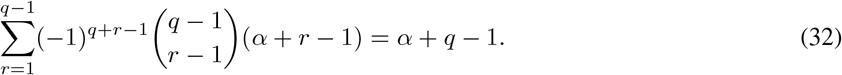

This equation, together with Eq. (30), implies that

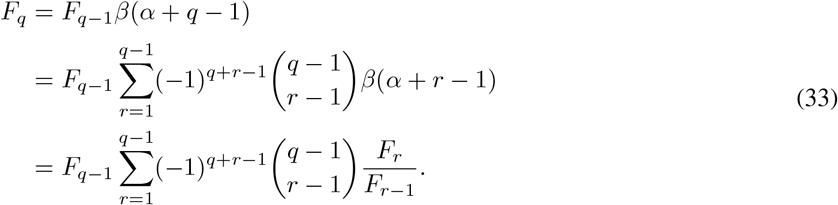

This equation expresses the *q*th-order factorial moment *F*_*q*_ in terms of lower-order factorial moments. In particular, setting *q* = 3 yields

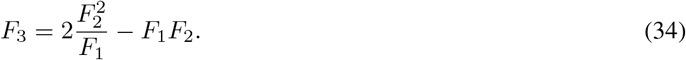

Since *F*_1_ = *(n), F*_2_ = *(n*^2^*) ™ (n)*, and *F*_3_ = *(n*^3^*) ™* 3*(n*^2^) + 2*(n)*, the third moment can be approximated in terms of the mean and variance as

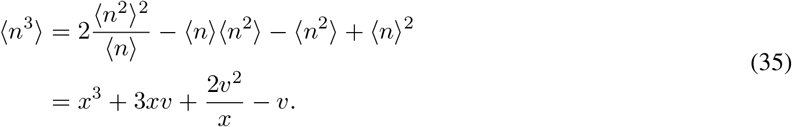

Substituting this expression into Eq. (6) yields the following MA system for the negative-binomial closure:

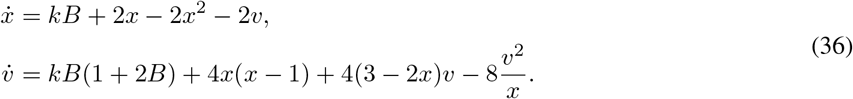

Note that Eq. (35) gives the second-order negative-binomial closure scheme. Higher-order closure schemes can be systematically derived from Eq. (33), which expresses the *q*th-order factorial moment *F*_*q*_ in terms of the lower-order factorial moments *F*_1_, …, *F*_*q™*1_. We emphasize that, although the second-order negative-binomial closure was previously proposed in Ref. [33], its extension to higher orders is introduced here for the first time.

The following theorem characterizes the stability of the negative-binomial closure (see Appendix C for the proof).

The results are essentially the same as those obtained for the log-normal closure.

### Theorem 4.

For all choices of *B* and *k*, the negative-binomial MA system given in Eq. (36) possesses a unique physically accessible fixed point, which is locally asymptotically stable. Moreover, the negative-binomial closure is stable if and only if

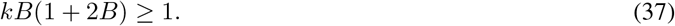

In other words, when *kB*(1 + 2*B*) ≥ 1, for any initial condition in *D*_phys_, the trajectory (*x*(*τ*), *v*(*τ*)) of the MA system remains in *D*_phys_ for all *τ* ≥ 0 and converges to the unique physically accessible fixed point.

Combining Theorems 2 and 4, we see that both the log-normal and negative-binomial closures have a unique physically accessible fixed point and are stable under the same mild condition. Fig. 4 illustrates the typical phase portraits of the negative-binomial MA system for *kB*(1 + 2*B*) *>* 1 and *kB*(1 + 2*B*) *<* 1, respectively. In the former case, all trajectories starting in *D*_phys_ remain within *D*_phys_ and converge to the unique physically accessible fixed point (Fig. 4(a)). In the latter case, some trajectories still remain within *D*_phys_ and converge to the unique physically accessible fixed point (green region in Fig. 4(b)); however, other trajectories enter the negative-variance region and thus become physically meaningless (orange region in Fig. 4(b)).

**FIG. 4.**
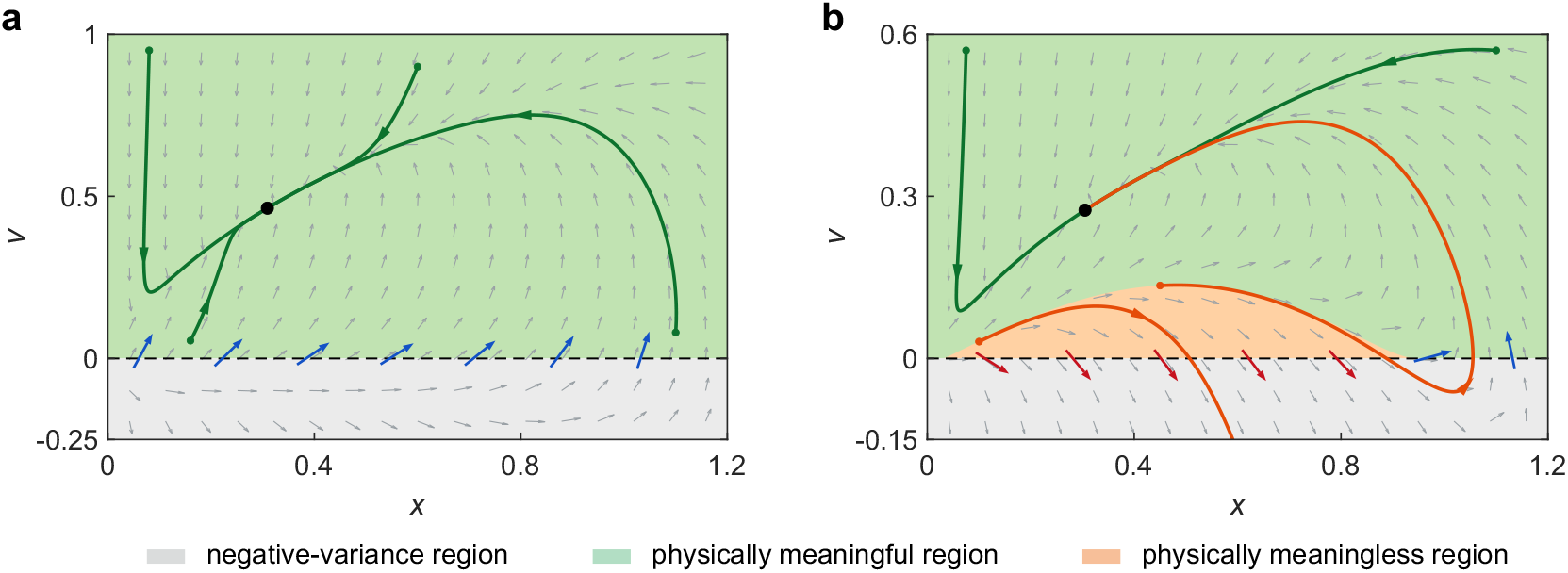
Typical phase portraits of the MA system for the negative-binomial closure. **(a)** The case *kB*(1 + 2*B*) *>* 1. The model parameters are chosen as *B* = 1.5, *k* = 1*/*3, and *kB*(1 + 2*B*) = 2. **(b)** The case *kB*(1 + 2*B*) *<* 1. The model parameters are chosen as *B* = 0.3, *k* = 5*/*12, and *kB*(1 + 2*B*) = 0.2. In (a) and (b), the grey region represents the negative-variance region. The green region indicates the set of initial conditions giving rise to physically meaningful trajectories (these trajectories remain within the physically meaningful region *D*_phys_ for all times and converge to the unique physically accessible fixed point). The orange region indicates the set of initial conditions giving rise to physically meaningless trajectories (these trajectories enter the negative-variance region at some time points).

## 6. Accuracy of the three moment-closure methods

Thus far, we have compared the stability of three distinct moment-closure methods for a bursty gene expression model with nonlinear degradation. Our results show that the log-normal and negative-binomial closures are substantially more stable than the Gaussian closure — the Gaussian closure is unstable for all parameter sets, whereas the log-normal and negative-binomial closures are stable under mild conditions.

In addition to stability, the performance of a moment-closure method is also evaluated in terms of its accuracy, which characterizes how closely the approximate moments predicted by the closure method agree with the true moments of the underlying stochastic dynamics. The accuracy of a moment-closure method is typically quantified by the relative errors between the approximate and true moments. Specifically, let *x*_true_(*τ*) and *v*_true_(*τ*) denote the true mean and variance of protein numbers at time *τ* for the original stochastic model, and let *x*_approx_(*τ*) and *v*_approx_(*τ*) denote the approximate mean and variance predicted by a particular moment-closure method. The time-averaged relative errors of the mean and variance are then defined, respectively, by

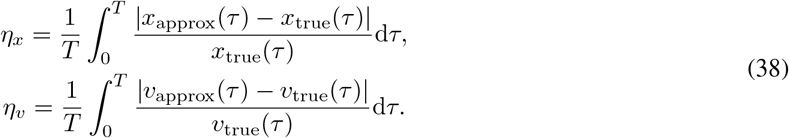

Here the termination time *T* is chosen such that the original stochastic system has nearly reached its steady state; specifically, *T* is defined as the time at which the Kullback–Leibler divergence between the time-dependent protein number distribution and the steady-state distribution becomes less than 10^*™*3^.

To further evaluate the accuracy of the three moment-closure methods, we randomly select 120 different parameter sets for the stochastic model, covering a broad region of the parameter space. The parameter sets are chosen such that both the log-normal and negative-binomial closures are stable. Since the Gaussian moment-closure method is inherently unstable, we restrict the parameter sets to those for which the Gaussian MA system possesses at least one physically accessible fixed point that is locally asymptotically stable. For each parameter set, the initial protein number is assumed to follow a Poisson distribution with a randomly selected mean *λ*_0_. For the Gaussian closure, only convergent trajectories of the MA system are retained, whereas divergent trajectories are excluded from the accuracy analysis.

Fig. 5(a),(b) show the time-averaged relative errors of the mean and variance as functions of *kB*(1 + 2*B*) for all parameter sets, while Fig. 5(c) shows the corresponding box plots of the relative errors. Clearly, among the three moment-closure methods, the negative-binomial closure is the most accurate for almost all parameter sets, whereas the Gaussian closure is the least accurate. The accuracy of the log-normal closure lies between those of the other two moment-closure methods. Interestingly, we also find that the relative errors of the Gaussian and negative-binomial closures increase slightly with *kB*(1 + 2*B*) (red circles and green triangles in Fig. 5(a),(b)), whereas those of the log-normal closure remain nearly unchanged as *kB*(1 + 2*B*) varies (blue squares in Fig. 5(a),(b)). Taking both stability and accuracy into account, the negative-binomial closure exhibits the best overall performance among the three moment-closure methods for the bursty gene expression model considered here.

**FIG. 5.**
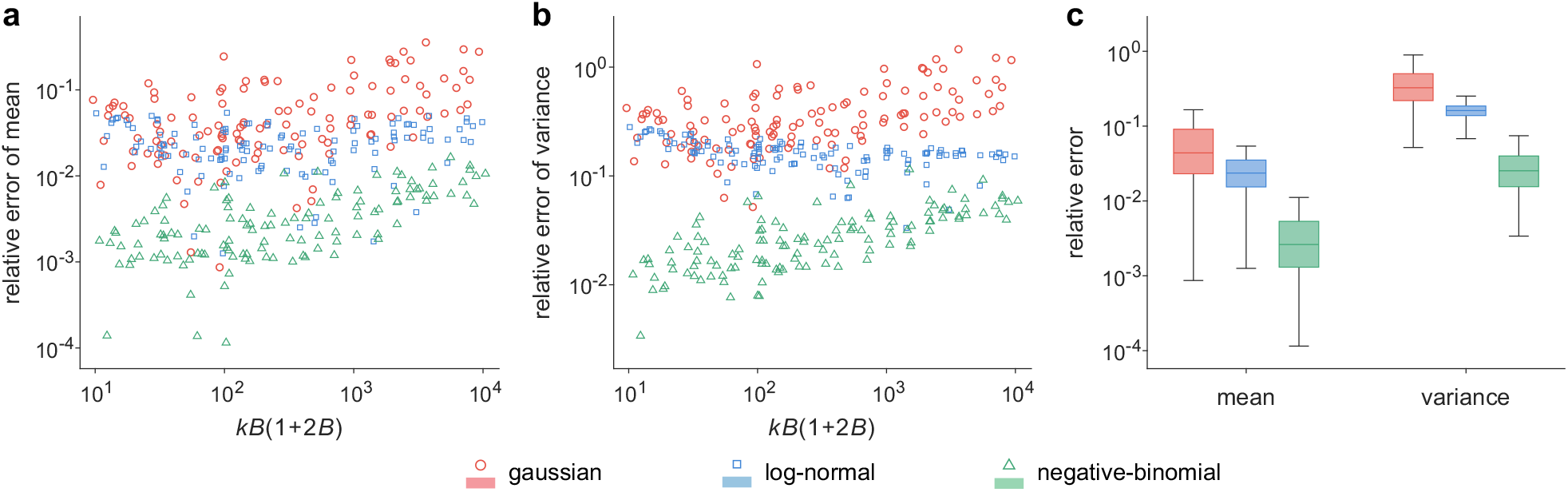
Accuracy of the Gaussian, log-normal, and negative-binomial closures. **(a)** Time-averaged relative errors of the mean for 120 randomly selected parameter sets as a function of *kB*(1 + 2*B*). **(b)** Time-averaged relative errors of the variance for all parameter sets as a function of *kB*(1 + 2*B*). In (a) and (b), red circles, blue squares, and green triangles represent the Gaussian, log-normal, and negative-binomial closures, respectively. The parameters are randomly sampled according to log_10_ *B ~ U* [*™*1, 1] and log_10_ *k~ U* [™4, 2], subject to the constraint log_10_(*kB*(1 + 2*B*)) ∈ [1, 4]. Here *U* [*a, b*] denotes the uniform distribution on the interval [*a, b*]. The initial protein number is assumed to follow a Poisson distribution with mean *λ*_0_, where *λ*_0_ is randomly sampled according to log_10_ *λ*_0_ *~ U* [*™*1, 1]. The true mean and variance are obtained by numerically solving the CME using the finite-state projection algorithm [19]. **(c)** Box plots of the relative errors of the mean and variance for the 120 parameter sets. The boxes extend from the first to the third quartiles, the center lines indicate the medians, and the whiskers extend to the most extreme data points within 1.5 times the interquartile range from the box edges.

So far, we have focused exclusively on second-order moment-closure approximations, in which the third-order moment is expressed in terms of the first- and second-order moments. In fact, all three moment-closure methods can be extended to higher-order approximations. For example, for the negative-binomial closure, the *q*th-order factorial moment *F*_*q*_ can be expressed in terms of the lower-order factorial moments *F*_1_, …, *F*_*q™*1_ based on Eq. (33). This yields a closed MA system of dimension *q ™* 1, corresponding to a (*q ™* 1)th-order approximation. To illustrate the performance of higher-order negative-binomial closures, we compare the accuracy of the second-to fifth-order approximations for the same 120 parameter sets considered above. For each approximation, we calculate the time-averaged relative errors of all retained factorial moments. The median relative errors across all parameter sets are summarized in Table 2. As the approximation order increases, the relative errors decrease substantially. In particular, the median relative error of the first-order (factorial) moment decreases from 0.262% to 0.002%, while that of the second-order factorial moment decreases from 0.410% to 0.005%. Under the fifth-order closure, the median relative errors of all five retained factorial moments are below 0.1%. These results further demonstrate the high accuracy of the negative-binomial moment-closure method.

**Table 2.** Median time-averaged relative errors of the retained factorial moments for the second-to fifth-order negative-binomial moment-closure approximations across 120 randomly selected parameter sets. “NA” in the table denotes “not available”.

| factorial moment | 2nd-order closure | 3rd-order closure | 4th-order closure | 5th-order closure |
| --- | --- | --- | --- | --- |
| first-order | 0.262% | 0.031% | 0.007% | 0.002% |
| second-order | 0.410% | 0.059% | 0.016% | 0.005% |
| third-order | NA | 0.124% | 0.027% | 0.009% |
| fourth-order | NA | NA | 0.077% | 0.018% |
| fifth-order | NA | NA | NA | 0.055% |

## 7 Conclusions and discussion

Moment-closure approximations are widely used to reduce the infinite hierarchy of moment equations arising in stochastic biochemical reaction networks to a finite-dimensional dynamical system. Their performance, however, cannot be assessed solely by comparing the approximate moments with those obtained from the CME. An equally fundamental requirement is that the closed moment equations generate physically meaningful dynamics. In particular, the predicted means and variances should remain nonnegative, trajectories should not diverge in parameter regimes where the underlying stochastic process is ergodic, and the long-time dynamics should approach a physically accessible fixed point. In this work, we have systematically investigated these issues for a bursty gene expression model with nonlinear protein degradation and compared three representative second-order closure schemes: the Gaussian, log-normal, and negative-binomial closures.

Our analysis reveals striking qualitative differences among these closure methods. For the Gaussian closure, we proved that the resulting MA system is unstable for all choices of the model parameters. Depending on the mean burst size *B* and the dimensionless protein synthesis rate *k*, the system may possess no physically accessible fixed point, a single unstable physically accessible fixed point, or two physically accessible fixed points with different stability properties. Consequently, the Gaussian closure necessarily admits initial conditions for which the predicted mean or variance becomes physically meaningless or the trajectory diverges. This result illustrates an important point: even a standard and widely used moment-closure approximation may fundamentally alter the global dynamical structure of the original stochastic system. Since the underlying stochastic gene expression model considered here is ergodic and converges to a unique stationary distribution, the additional fixed points, unstable fixed points, and divergent trajectories generated by the Gaussian closure are artifacts introduced by the closure approximation rather than genuine properties of the stochastic dynamics.

The situation is fundamentally different for the log-normal and negative-binomial closures. For both methods, we proved that the corresponding MA system possesses a unique physically accessible fixed point that is locally asymptotically stable for all parameter values. More importantly, both closures satisfy exactly the same global stability condition *kB*(1 + 2*B*) ≥ 1. When this condition holds, the physically meaningful region *D*_phys_ is forward invariant, and every trajectory starting from a physically admissible initial condition converges to the unique physically accessible fixed point. When this condition is violated, the physically accessible fixed point remains locally stable, but some trajectories can cross the boundary *v* = 0 and enter the negative-variance region. Thus, the existence and even local stability of a physically meaningful equilibrium alone are insufficient to guarantee the stability of a moment-closure approximation; the global geometry of the vector field, particularly its behavior on the boundary of the physically admissible region, must also be taken into account.

An important feature of this stability condition is that it has a direct biological interpretation. We showed that if the steady-state mean protein number of the original stochastic model exceeds one, then *kB*(1 + 2*B*) *>* 1 is automatically satisfied. Therefore, the stability condition for the log-normal and negative-binomial closures is expected to hold in a broad range of biologically relevant situations. Indeed, experimentally measured protein abundances are typically considerably greater than one molecule per cell across bacteria, yeast, and mammalian cells. This observation suggests that the instability occurring below the threshold *kB*(1 + 2*B*) = 1, although mathematically important for establishing the exact stability condition, may have limited practical relevance in many biological settings where protein copy numbers are not extremely low.

The negative-binomial distribution is particularly natural for gene expression because, unlike Gaussian and log-normal distributions, it is supported on nonnegative integers and assigns a nonzero probability to zero molecules. This structural compatibility with molecular copy numbers may partly explain the favorable performance observed here. In our numerical comparison over 120 randomly selected parameter sets, the negative-binomial closure gives the smallest errors in both the mean and variance for almost all cases considered, whereas the Gaussian closure exhibits the largest errors and the log-normal closure shows intermediate accuracy. Notably, divergent Gaussian trajectories were already excluded from this accuracy comparison, indicating that its inferior numerical performance persists even when attention is restricted to its convergent trajectories. We further developed higher-order negative-binomial closure schemes based on combinatorial identities between factorial moments of the negative-binomial distribution and demonstrated that increasing the closure order substantially improves predictive accuracy. Taking both stability and accuracy into account, the negative-binomial closure therefore exhibits the best overall performance among the three moment-closure methods for the bursty gene expression model studied here.

Several questions warrant further investigation. The present analysis is restricted to a one-species gene expression model with bursty synthesis, dimerization-mediated degradation, and second-order moment closures. It will be important to determine to what extent the conclusions obtained here extend to more complex biochemical reaction networks involving promoter switching [6, 7], feedback regulation [52, 53], multiple interacting molecular species [28, 29], or higher-order nonlinear reaction mechanisms [23, 32]. Another natural direction is to investigate higher-order extensions of the three moment-closure methods and determine whether their stability properties persist as additional moments are retained. More generally, it would be interesting to develop systematic principles for constructing closure functions that preserve intrinsic structural properties of the underlying stochastic process, such as nonnegativity, discreteness, and ergodicity. Such developments may lead to moment-closure schemes that are not only quantitatively accurate but also dynamically and physically consistent.

## Appendices

### A. Ergodicity of the original stochastic model

We now prove that the original stochastic Markovian model is ergodic. Specifically, let *X*_*t*_ denote the protein number at time *t*. Clearly, *X*_*t*_ is a continuous-time Markov chain on the state space *S* = *{*0, 1, 2, *…}* with transition rates

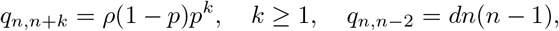

where the transition from microstate *n* to *n* + *k* is due to the synthesis reaction *G → G* + *kP*, whereas the transition from microstate *n* to *n ™* 2 is due to the degradation reaction 2*P →* ∅. Since the transitions *n → n* + 1 and *n → n ™* 2 have positive rates whenever they are admissible, it is easy to see that the process *X*_*t*_ is irreducible.

For any nonnegative function *V* : *S →* [0, *∞*), we define

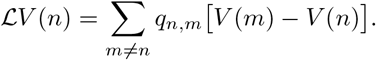

Here the operator *L* is referred to as the generator of the Markov process. Recall that the function *V* is said to be norm-like if its sublevel set {*n* ∈ *S* : *V* (*n*) *≤ K}* is finite for any *K* ≥ 0. To proceed, we need the following two lemmas, whose proofs can be found in [41].

#### Lemma 1

(Lyapunov criterion for non-explosivity). Let *X*_*t*_ be a continuous-time Markov chain on a countable state space *S* with generator *L*. Suppose there exists a norm-like function *V* : *S →* [0, *∞*) and a constant *c* ≥ 0 such that

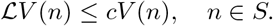

Then the process *X*_*t*_ is non-explosive and is therefore well-defined for all *t* ≥ 0.

#### Lemma 2

(Foster-Lyapunov criterion for positive recurrence). Let *X*_*t*_ be an irreducible, non-explosive continuous-time Markov chain on a countable state space *S* with generator *L*. Suppose there exist a function *V* : *S →* [0, *∞*) and a finite set *K ⊂ S* such that

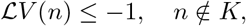

while sup*n*∈*K* | *LV* (*n*)| *< ∞*. Then the process *X*_*t*_ is positive recurrent.

We next apply these two lemmas to prove that the process *X*_*t*_ is non-explosive and positive recurrent. Applying the generator to the norm-like function *V* (*n*) = *n* + 1 yields

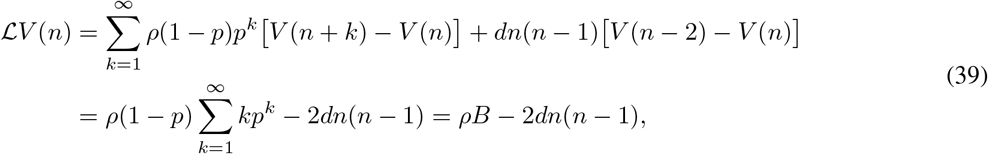

where *B* = *p/*(1 *™ p*) is the mean protein burst size. It follows immediately that

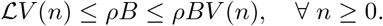

Hence, the Lyapunov criterion in Lemma 1 implies that the process is non-explosive. Moreover, since the negative quadratic term in Eq. (39) dominates for sufficiently large *n*, there exists *K >* 0 such that

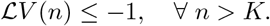

The Foster-Lyapunov criterion in Lemma 2 therefore implies that the process is positive recurrent [41]. Since the process is also irreducible, it admits a unique stationary distribution and is ergodic. In particular, regardless of the initial condition, the time-dependent distribution converges to the stationary distribution as 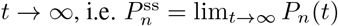.

### B. Proof of two combinatorial identities

Here we prove the two combinatorial identities given in Eq. (31). For any *q* ≥ 2, it follows from the binomial expansion that

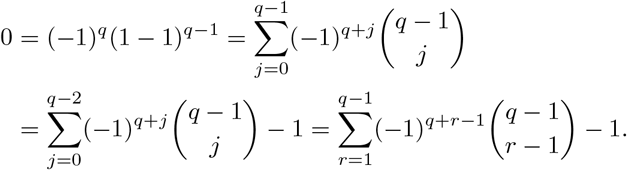

This proves the first identity in Eq. (31). On the other hand, for any *q* ≥ 3, applying the binomial expansion together with the combinatorial identity

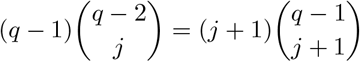

yields

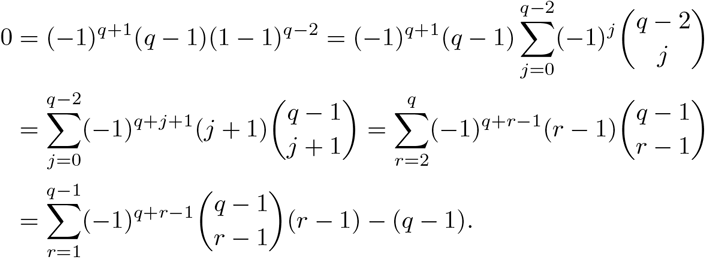

This proves the second identity in Eq. (31).

### C. Proof of Theorem 4

We now prove Theorem 4. Recall that the MA system associated with the negative-binomial closure is

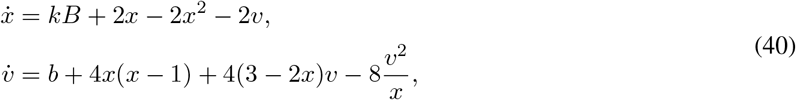

where, for convenience, we set *b* = *kB*(1 + 2*B*). The proof consists of two steps. We first establish the existence and uniqueness of a physically accessible fixed point. We then prove that this fixed point is locally asymptotically stable for all *B, k >* 0 and determine the exact condition under which it is globally asymptotically stable in *D*_phys_.

#### Lemma 3

(Uniqueness and physical accessibility). For every *B, k >* 0, the negative-binomial closure has exactly one physically accessible fixed point in *D*_phys_.

*Proof*. At equilibrium, the first equation of Eq. (40) gives

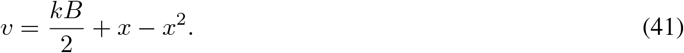

Substituting this expression into the second equation of Eq. (40), we find that the equilibrium condition reduces to the scalar quadratic equation

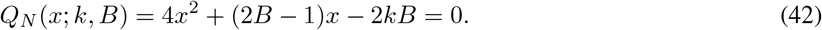

The product of its two roots is *™kB/*2 *<* 0. Hence, the two roots are both real and have opposite signs. It then follows that Eq. (42) has exactly one positive root, which we denote by *x*_*∗*_. At this root, Eq. (42) gives

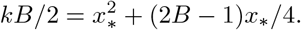

Substituting this relation into Eq. (41) yields

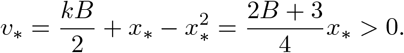

Therefore, (*x*_*∗*_, *v*_*∗*_) ∈ *D*_phys_. Since Eq. (42) has no other positive root, this is the unique physically accessible fixed point of the negative-binomial MA system.

#### Lemma 4

(Local and global asymptotic stability). For every *B, k >* 0, the unique physically accessible fixed point of the negative-binomial closure is locally asymptotically stable. Moreover, it is globally asymptotically stable in *D*_phys_ if and only if *b* = *kB*(1 + 2*B*) ≥ 1.

*Proof*. We first establish local asymptotic stability. For a real 2 *×* 2 Jacobian matrix *J*, both eigenvalues have negative real parts if and only if tr*J <* 0 and det *J >* 0. At an equilibrium satisfying Eq. (41), direct calculation gives

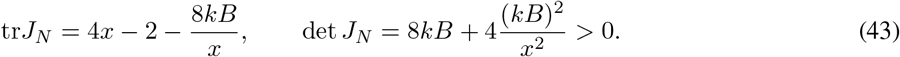

At the unique physically accessible fixed point (*x*_*∗*_, *v*_*∗*_), Eq. (42) further gives

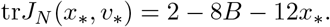

In addition, the same equation implies that *x*_*∗*_(4*x*_*∗*_ + 2*B ™* 1) = 2*kB >* 0, and hence 4*x*_*∗*_ + 2*B >* 1. Since *x*_*∗*_ *>* 0 and *B >* 0, we further have 6*x*_*∗*_ + 4*B >* 1, so that

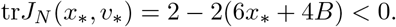

Together with Eq. (43), this proves that the unique physically accessible fixed point is locally asymptotically stable for all *B, k >* 0.

We next characterize its global stability. We first demonstrate that the condition *b* ≥ 1 is necessary. On the boundary *v* = 0, the second equation of Eq. (40) reduces to

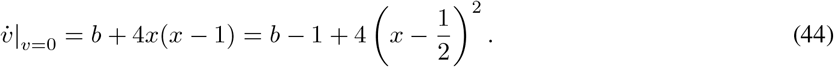

If *b <* 1, the right-hand side is negative for *x* in an open interval containing 1*/*2. Therefore, by continuity of the vector field, there exist initial conditions in *D*_phys_ sufficiently close to this boundary segment for which the corresponding trajectories cross the boundary *v* = 0 and enter the negative-variance region *v <* 0. Thus, *D*_phys_ is not forward invariant when *b <* 1, and global stability in *D*_phys_ is impossible.

It remains to prove that *b* ≥ 1 is sufficient. Assume henceforth that *b* ≥ 1. Let (*x*(*τ*), *v*(*τ*)) be a trajectory starting in the interior of *D*_phys_, and let *τ*_e_ denote its first possible exit time from the interior of *D*_phys_. We first derive bounds on the trajectory for 0 *≤ τ < τ*_e_. Since *v >* 0, the first equation of Eq. (40) gives 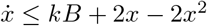. Let

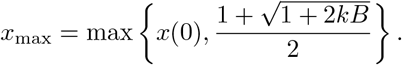

The right-hand side of the above inequality is nonpositive at *x* = *x*_max_ and strictly negative for *x > x*_max_. It follows that 0 *< x*(*τ*) *≤ x*_max_ for 0 *≤ τ < τ*_e_. Using this bound in the second equation of Eq. (40), we obtain

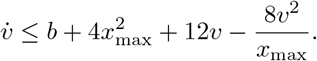

Since the right-hand side tends to *™∞* as *v → ∞*, a standard comparison argument indicates that there exists a constant *v*_max_ *>* 0 such that 0 *< v*(*τ*) *≤ v*_max_ for 0 *≤ τ < τ*_e_. Therefore, finite-time blow-up cannot occur before the trajectory reaches the boundary of *D*_phys_.

We next show that the boundary *v* = 0 cannot be reached from the interior. If *b >* 1, then Eq. (44) gives 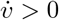 for every *x >* 0, and hence no trajectory starting with *v >* 0 can cross the boundary *v* = 0. The borderline case *b* = 1 requires slightly more care. In this case, the right-hand side of Eq. (44) vanishes only at *Q* = (1*/*2, 0). Let

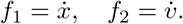

At *Q*, we have *f*_1_(*Q*) = *kB* + 1*/*2 *>* 0, *f*_2_(*Q*) = 0, *∂*_*x*_*f*_2_(*Q*) = 0, and *∂*_*xx*_*f*_2_(*Q*) = 8. Consider the solution passing through *Q* at time *τ*_0_, i.e. (*x*(*τ*_0_), *v*(*τ*_0_)) = *Q*. Along this solution, we have

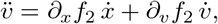

so that 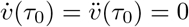. Differentiating once more and evaluating at *τ*_0_ gives 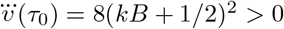 Taylor expansion around *τ*_0_ therefore yields *>* 0.

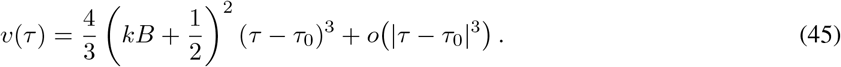

Hence, the solution through *Q* lies in the region *v <* 0 for *τ < τ*_0_ sufficiently close to *τ*_0_. If a trajectory starting in the interior *v >* 0 were to reach *Q*, uniqueness of solutions would force it to coincide locally with this trajectory, which is impossible. At every other point on the boundary *v* = 0, Eq. (44) gives 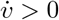. We conclude that the boundary *v* = 0 cannot be reached from the interior of *D*_phys_ for any *b* ≥ 1.

It remains to exclude approach to the singular boundary *x* = 0. To this end, we choose 0 *< δ <* min*{x*(0), 1*}* and define *K* = *v*_max_*/δ*. By decreasing *δ* if necessary, we may assume that *K* ≥ max*{*1, *b/*(*kB*)*}*. Suppose that the trajectory enters the strip 0 *< x ≤ δ* through the line *x* = *δ*. Since *v ≤ v*_max_ = *Kδ* at the entry point, the trajectory initially lies below the line *v* = *Kx*. On this line, direct substitution into Eq. (40) gives

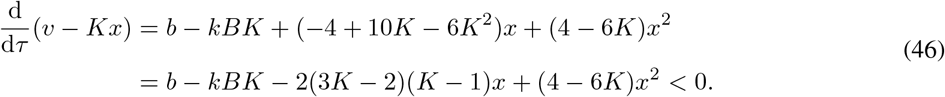

Here the last inequality follows from *K* ≥ 1, *K* ≥ *b/*(*kB*), and *x >* 0. Therefore, the vector field along *v* = *Kx* points toward the region *v < Kx*, and the trajectory cannot cross this line from below. Consequently, *v*(*τ*) *≤ Kx*(*τ*) whenever 0 *< x*(*τ*) *≤ δ*. The first equation of Eq. (40) then gives

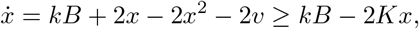

where we have used 0 *< x ≤ δ <* 1. Let

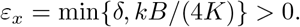

Whenever 0 *< x ≤ ε*_*x*_, we have 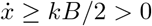. Therefore, a trajectory cannot cross the line *x* = *ε*_*x*_ from right to left, and in particular it cannot approach the singular boundary *x* = 0.

We can now conclude that the trajectory exists globally and remains in the physically meaningful region. Indeed, if *τ*_e_ *< ∞*, then the bounds obtained above imply that the trajectory remains in a compact subset of *{*(*x, v*) : *x >* 0*}* whenever 0 *≤ τ < τ*_e_. Since the vector field is continuously differentiable on this set, the solution can be extended beyond *τ*_e_. On the other hand, the preceding arguments show that neither *v* = 0 nor *x* = 0 can be reached at *τ*_e_, which is a contradiction. Hence *τ*_e_ = *∞*. Moreover,

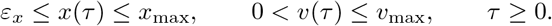

Thus, every positive orbit is bounded and remains a positive distance away from the singular boundary *x* = 0.

It remains only to establish convergence to the unique physically accessible fixed point. Let ***f*** = (*f*_1_, *f*_2_)^T^ denote the vector field in Eq. (40). The interior of *D*_phys_ is simply connected. Consider the Dulac function

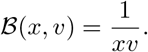

A direct calculation gives

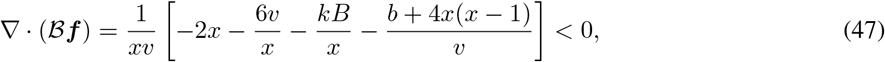

where we have used *b* + 4*x*(*x ™* 1) = *b ™* 1 + 4(*x ™* 1*/*2)^2^ ≥ 0. Thus, by the Bendixson–Dulac criterion, the system admits no periodic orbit lying entirely in the interior of *D*_phys_ [54].

Let *ω*(*x*(0), *v*(0)) denote the omega-limit set of a positive orbit. Since the orbit is bounded and remains away from *x* = 0, its omega-limit set is nonempty and compact. Suppose that *ω*(*x*(0), *v*(0)) contains no fixed point. The Poincaré–Bendixson theorem then implies that the omega-limit set must be a periodic orbit [55]. Such a periodic orbit cannot intersect the boundary *v* = 0. Indeed, away from *Q*, we have 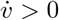 on *v* = 0, so its backward orbit immediately enters *v <* 0; at *Q*, the same conclusion follows from Eq. (45). Hence, any such periodic orbit would have to lie entirely in the interior of *D*_phys_, contradicting the Bendixson–Dulac criterion.

Therefore, *ω*(*x*(0), *v*(0)) must contain a fixed point. According to Lemma 3, the only physically accessible fixed point is (*x*_*∗*_, *v*_*∗*_), and hence (*x*_*∗*_, *v*_*∗*_) ∈ *ω*(*x*(0), *v*(0)). Since (*x*_*∗*_, *v*_*∗*_) is locally asymptotically stable, it possesses an attracting neighborhood. Because it belongs to the omega-limit set, the trajectory must eventually enter this neighborhood, after which it converges to (*x*_*∗*_, *v*_*∗*_). Hence (*x*(*τ*), *v*(*τ*)) *→* (*x*_*∗*_, *v*_*∗*_) as *τ → ∞*.

Thus, when *b* ≥ 1, every positive trajectory remains in *D*_phys_ for all forward time and converges to the unique physically accessible fixed point. Combining the necessity and sufficiency arguments, we conclude that the unique physically accessible fixed point is globally asymptotically stable in *D*_phys_ if and only if *b* = *kB*(1 + 2*B*) ≥ 1.

Lemmas 3 and 4 complete the proof of Theorem 4.

## Acknowledgements

C. J. acknowledges support from National Natural Science Foundation of China with grant Nos. 12526215, T2541031, and 12271020.

