## Supplementary Material for "Dynamical stability and accuracy of moment closure approximations for bursty gene expression with nonlinear degradation"

#### Contents

**1 Proof of Proposition 1** **2**

**2 Proof of Theorem 2** **6**

### 1 Proof of Proposition 1

We now prove Proposition 1 in the main text. Recall that the Gaussian closure, $\langle n^3\rangle = 3\langle n^2\rangle\langle n\rangle - 2\langle n\rangle^3$, gives the closed MA system

$$\dot{x} = kB + 2x - 2x^2 - 2v,$$
$$\dot{v} = kB(1+2B) + 4x(x-1) + 8(1-x)v.$$

At equilibrium, the first equation gives $v = kB/2 + x - x^2$. Substituting this expression into the second equation yields

$$Q_G(x; k, B) = 8x^3 - 12x^2 + 4(1-kB)x + kB(5+2B) = 0.$$

For later use, define

$$R_B(k) = 32B^3k^3 - \left(108B^4 + 324B^3 + 195B^2\right)k^2 + 24Bk + 4.$$

The Jacobian matrix is

$$J_G(x, v) = \begin{pmatrix} 2-4x & -2 \\ 8x-4-8v & 8-8x \end{pmatrix}.$$

At an equilibrium, direct calculation gives $\det J_G = 8(6x^2 - 6x + 1 - kB)$ and $\operatorname{tr} J_G = 10 - 12x$. Since

$$\partial_x Q_G = 4(6x^2 - 6x + 1 - kB),$$

every equilibrium satisfies

$$\det J_G = 2\partial_x Q_G. \tag{1}$$

This identity provides a direct connection between the ordering and multiplicity of the roots of $Q_G$ and the local stability of the corresponding equilibria. Because the proof of the proposition is somewhat lengthy, we divide it into three lemmas. The first determines the roots of $Q_G$ and identifies the physically accessible equilibria, the second considers the two boundary values $k_1(B)$ and $k_2(B)$, and the third determines the stability of the two equilibria away from these boundaries.

**Lemma 1** (Roots and physically accessible equilibria)**.** For every $B > 0$, $R_B(k) = 0$ has one negative root and two positive roots $k_1(B) < k_2(B)$, satisfying $0 < k_1(B) < 1/B < k_2(B)$. If $0 < k < k_1(B)$ or $k > k_2(B)$, the Gaussian system has exactly two physically accessible fixed points. If $k_1(B) < k < k_2(B)$, it has no physically accessible fixed point. At $k = k_1(B)$ or $k = k_2(B)$, it has exactly one physically accessible fixed point, corresponding to a repeated positive root of $Q_G$.

*Proof.* The discriminant of $Q_G$, regarded as a cubic polynomial in $x$, is $\Delta_G(k, B) = 64R_B(k)$. Thus, the sign of $R_B(k)$ determines whether $Q_G$ has three distinct real roots, a repeated root, or one real root together with a complex-conjugate pair [1].

The coefficient signs of $R_B(k)$ are $+, -, +, +$, so Descartes' rule of signs shows that $R_B$ has at most two positive roots. Moreover, it is clear that

$$R_B(0) = 4 > 0, \quad R_B(1/B) = 60 - (108B^2 + 324B + 195) < 0,$$

and $R_B(k) \to +\infty$ as $k \to +\infty$. Therefore, $R_B$ has one positive root in $(0, 1/B)$ and another in $(1/B, \infty)$, and these are its only positive roots. Similarly, the coefficient signs of $R_B(-k)$ are $-, -, -, +$. Descartes' rule of signs, together with the fact that $R_B(k) \to -\infty$ as $k \to -\infty$, shows that $R_B$ has exactly one negative root. Consequently, $R_B(k) > 0$ for $0 < k < k_1(B)$ or $k > k_2(B)$, whereas $R_B(k) < 0$ for $k_1(B) < k < k_2(B)$.

When $R_B(k) > 0$, the cubic $Q_G$ has three distinct real roots. Since $Q_G(-\infty; k, B) < 0 < Q_G(0; k, B)$, one of these roots is negative. Vieta's relations then imply that the remaining two roots are positive: their product is positive and their sum is positive. For any root $x$ of $Q_G$, the corresponding equilibrium variance can be written as

$$v = \frac{x(x-1)(2B+3)}{4x-(2B+5)}. \tag{2}$$

Let $x_T = (2B+5)/4$. For $1 \leq x \leq x_T$, we have

$$Q_G(x; k, B) = 4x(x-1)(2x-1) + kB(2B+5-4x) > 0.$$

Hence, each positive root must lie either in the interval $(0,1)$ or in the interval $(x_T, \infty)$. In either interval, the numerator and denominator in Eq. (2) have the same sign, and therefore both positive roots correspond to $v > 0$. In contrast, the negative root gives $v < 0$ and is thus physically inaccessible.

When $R_B(k) < 0$, the cubic $Q_G$ has only one real root. Since $Q_G(-\infty; k, B) < 0 < Q_G(0; k, B)$, this root is negative, and hence no physically accessible fixed point exists. Finally, when $k = k_1(B)$ or $k = k_2(B)$, the two positive roots coalesce into a repeated positive root. The strict positivity of $Q_G$ on $[1, x_T]$, together with Eq. (2), shows that the corresponding variance remains positive. Thus, in either case the system has exactly one physically accessible fixed point. □

**Lemma 2** (The two boundary values)**.** At $k = k_1(B)$ or $k = k_2(B)$, the two physically accessible fixed points coalesce into a single unstable saddle-node fixed point. The saddle-node is nondegenerate except at $(B, k) = (5/18, 3/5)$, where the Jacobian has a double zero eigenvalue and the saddle-node is degenerate.

*Proof.* Let $k_s \in \{k_1(B), k_2(B)\}$ and let $x_s$ denote the repeated positive root. Then

$$Q_G(x_s; k_s, B) = 0, \quad \partial_x Q_G(x_s; k_s, B) = 0,$$

so Eq. (1) gives $\det J_G = 0$. Moreover, since $Q_G(1/2; k, B) = kB(2B+3) > 0$, it is easy to see that

$$\partial_{xx} Q_G(x_s; k_s, B) = 24(2x_s - 1) \neq 0.$$

We also have

$$\partial_k Q_G(x_s; k_s, B) = B(2B+5-4x_s) \neq 0.$$

Indeed, if $x_T = (2B+5)/4$, then

$$Q_G(x_T; k, B) = 4x_T(x_T - 1)(2x_T - 1) > 0,$$

so the repeated root cannot coincide with $x_T$. Thus, the repeated root is a quadratic fold with respect to the parameter $k$.

If $x_s \neq 5/6$, then $\operatorname{tr} J_G = 10 - 12x_s \neq 0$, so the zero eigenvalue is simple. Since $\partial\dot{x}/\partial v = -2 \neq 0$, the equilibrium condition can be reduced locally to the scalar equation $Q_G = 0$. Together with $\partial_{xx} Q_G \neq 0$ and $\partial_k Q_G \neq 0$, these conditions imply a nondegenerate saddle-node bifurcation [2, 3]. Equivalently, the reduced dynamics on the one-dimensional center manifold has a nonzero quadratic leading term, and the saddle-node fixed point is therefore unstable.

The exceptional case $x_s = 5/6$ occurs only at $B = 5/18$, $k = 3/5$, and $(x_s, v_s) = (5/6, 2/9)$. At this point, the Jacobian is nonzero and nilpotent, with two zero eigenvalues. Together with $\partial_{xx} Q_G \neq 0$ and $\partial_k Q_G \neq 0$, the standard nilpotent saddle-node criterion identifies a degenerate saddle-node. The corresponding normal form contains an outward sector, and hence the fixed point is unstable [2, 3]. □

**Lemma 3** (Stability of the two fixed points)**.** Whenever the Gaussian system has two physically accessible fixed points, the smaller-mean fixed point is a saddle. The larger-mean fixed point has the following stability properties.

(i) If $0 < B \leq 5/18$, it is locally asymptotically stable for $k \in (0, k_1(B)) \cup (k_2(B), \infty)$.

(ii) If $B > 5/18$, then $k_3(B) := 10/[9B(6B + 5)] < k_1(B) < k_2(B)$. The larger-mean fixed point is locally asymptotically stable for $k \in (0, k_3(B)) \cup (k_2(B), \infty)$, is an unstable weak focus at $k = k_3(B)$, and is an unstable node or focus for $k \in (k_3(B), k_1(B))$.

*Proof.* Let the two positive roots be $x_*^{(1)} < x_*^{(2)}$, in agreement with the notation used in the main text. Since they are, respectively, the middle and largest roots of the cubic $Q_G$, whose leading coefficient is positive, we have

$$\partial_x Q_G(x_*^{(1)}; k, B) < 0, \quad \partial_x Q_G(x_*^{(2)}; k, B) > 0.$$

It then follows from Eq. (1) that the smaller-mean fixed point has negative determinant and is therefore a saddle, whereas the larger-mean fixed point has positive determinant. Consequently, the stability of the latter is determined entirely by the sign of $\operatorname{tr} J_G = 10 - 12x_*^{(2)}$.

We first consider $k > k_2(B)$. By Lemma 1, $kB > 1$. For $0 < x \leq 1/2$, both terms in

$$Q_G(x; k, B) = 4x(x - 1)(2x - 1) + kB(2B + 5 - 4x)$$

are nonnegative, with the second term being strictly positive. For $1/2 < x < 1$, we have

$$4x(1 - x)(2x - 1) < 1, \quad kB(2B + 5 - 4x) > 1,$$

and hence $Q_G(x; k, B) > 0$ again. Together with the absence of roots in $[1, x_T]$, this yields

$$x_*^{(2)} > x_T > \frac{5}{6}.$$

Therefore $\operatorname{tr} J_G < 0$, and the larger-mean fixed point is locally asymptotically stable for every $k > k_2(B)$.

We next consider the lower two-fixed-point interval. A change in stability of the larger-mean fixed point can occur only when its trace vanishes, that is, when $x_*^{(2)} = 5/6$. Substituting $x = 5/6$ into $Q_G = 0$ gives

$$k = k_3(B) := \frac{10}{9B(6B + 5)}.$$

At this parameter value,

$$\det J_G = \frac{4(18B - 5)}{9(6B + 5)}. \tag{3}$$

If $0 < B < 5/18$, the determinant in Eq. (3) is negative. Hence the trace-zero root cannot correspond to the larger root, whose determinant is positive. If $B = 5/18$, then $k_3 = 3/5 = k_1(B)$, so the trace-zero point occurs only at the endpoint of the lower two-fixed-point interval. Since $x_*^{(2)} \to 1$ and $\operatorname{tr} J_G \to -2$ as $k \to 0^+$, and no trace-zero point is encountered in the interior of the interval, the larger-mean fixed point is locally asymptotically stable throughout $0 < k < k_1(B)$ for $0 < B \leq 5/18$.

Now suppose that $B > 5/18$. By Eq. (3), the determinant at $k = k_3(B)$ is positive, and

$$Q'_G\left(\frac{5}{6}; k_3, B\right) > 0.$$

Thus $Q_G(x; k_3, B) < 0$ for $x < 5/6$ sufficiently close to $5/6$. Since

$$Q_G\left(\frac{1}{2}; k_3, B\right) > 0, \quad Q_G\left(\frac{5}{6}; k_3, B\right) = 0,$$

the intermediate value theorem implies that $Q_G$ has another positive root in $(1/2, 5/6)$. Hence $k_3$ lies in a two-fixed-point parameter regime. Moreover,

$$k_3 B = \frac{10}{9(6B+5)} < 1,$$

so that $k_3 < 1/B < k_2(B)$. By Lemma 1, it follows that

$$0 < k_3(B) < k_1(B) < k_2(B).$$

Near $k = k_3(B)$, the implicit function theorem gives

$$\frac{\mathrm{d}x_*^{(2)}}{\mathrm{d}k} = -\frac{(Q_G)_k}{(Q_G)_x} < 0,$$

because

$$(Q_G)_x > 0, \quad (Q_G)_k = B(2B + 5 - 4x_*^{(2)}) > 0.$$

It follows that $\operatorname{tr} J_G = 10 - 12x_*^{(2)}$ increases through zero as $k$ increases through $k_3(B)$. Therefore, the larger-mean fixed point is locally asymptotically stable for $0 < k < k_3(B)$ and unstable for $k_3(B) < k < k_1(B)$. In the latter interval, both the trace and determinant are positive, so the fixed point is an unstable node or focus.

It remains to determine the nonlinear stability at the boundary $k = k_3(B)$, where the linearization has purely imaginary eigenvalues. Define

$$x_H = \frac{5}{6}, \quad v_H = \frac{k_3 B}{2} + \frac{5}{36}, \quad \omega^2 = \det J_G(x_H, v_H) = \frac{4(18B-5)}{9(6B+5)} > 0.$$

Introducing the coordinates

$$u = x - x_H, \quad y = \frac{\frac{4}{3}u + 2(v - v_H)}{\omega},$$

transforms the Gaussian system at $k = k_3(B)$ exactly into

$$\dot{u} = -\omega y - 2u^2,$$
$$\dot{y} = \omega u + \frac{16}{\omega}u^2 - 8uy.$$

The linearized system therefore has eigenvalues $\pm i\omega$.

Write the nonlinear terms as

$$F(u, y) = -2u^2, \quad G(u, y) = \frac{16}{\omega}u^2 - 8uy.$$

Their only nonzero second-order derivatives at the origin are

$$F_{uu} = -4, \quad G_{uu} = \frac{32}{\omega}, \quad G_{uy} = -8,$$

while all third-order derivatives vanish. For a planar system whose linear part is $\dot{u} = -\omega y$, $\dot{y} = \omega u$, the standard formula for the first Lyapunov coefficient therefore reduces to

$$\begin{aligned}\ell_1 &= \frac{1}{16\omega}\left[F_{uy}(F_{uu} + F_{yy}) - G_{uy}(G_{uu} + G_{yy}) - F_{uu}G_{uu} + F_{yy}G_{yy}\right] \\ &= \frac{24}{\omega^2} > 0.\end{aligned}$$

Thus the equilibrium at $k = k_3(B)$ is a weak focus of first order and, since $\ell_1 > 0$, it is unstable. This completes the nonlinear classification at the boundary, where the trace–determinant criterion alone is inconclusive. □

Lemmas 1-3 prove all statements in Proposition 1 in the main text.

### 2 Proof of Theorem 2

We now prove Theorem 2 in the main text. Recall that the log-normal closure, $\langle n^3 \rangle = \langle n^2 \rangle^3 / \langle n \rangle^3$, gives the closed MA system

$$\begin{aligned}\dot{x} &= kB + 2x - 2x^2 - 2v, \\ \dot{v} &= b + 4x^3 + 4x^2 - 4x + 4xv + 8v - 4\frac{(v + x^2)^3}{x^3},\end{aligned} \tag{4}$$

where $b = kB(1 + 2B)$. We divide the proof into two lemmas. The first establishes the existence and uniqueness of a physically accessible fixed point, while the second proves its local asymptotic stability and determines the exact condition for global asymptotic stability in $\mathcal{D}_{\text{phys}}$.

**Lemma 4** (Uniqueness and physical accessibility)**.** For every $B, k > 0$, the log-normal closure has exactly one physically accessible fixed point in $\mathcal{D}_{\text{phys}}$.

*Proof.* At equilibrium, the first equation of Eq. (4) gives

$$v = \frac{kB}{2} + x - x^2. \tag{5}$$

Substituting this expression into the second equation of Eq. (4) reduces the equilibrium condition to

$$\begin{aligned}Q_L(x; k, B) = (4kB + 8)x^4 &+ (4kB^2 + 10kB - 8)x^3 \\ &- 12kBx^2 - 6k^2B^2x - k^3B^3 = 0.\end{aligned} \tag{6}$$

For $B, k > 0$, the nonzero coefficient sequence of $Q_L$ has exactly one sign change: the leading coefficient is positive, the quadratic, linear, and constant coefficients are negative, and either sign of the cubic coefficient leaves exactly one sign change. Descartes' rule of signs therefore implies that $Q_L$ has at most one positive root. Since

$$Q_L(0; k, B) = -k^3B^3 < 0, \quad \lim_{x \to +\infty} Q_L(x; k, B) = \infty,$$

it follows that $Q_L$ has exactly one positive root, denoted by $x_*$.

It remains to show that the corresponding variance is positive. Let $x_0 = (1 + \sqrt{1 + 2kB})/2$, the positive root of $kB/2 + x - x^2 = 0$. Equation (5) shows that $v > 0$ precisely when $0 < x < x_0$. Using $x_0^2 = x_0 + kB/2$, direct substitution gives

$$Q_L(x_0; k, B) = 2kB(2B + 3)\left[\left(1 + \frac{kB}{2}\right)x_0 + \frac{kB}{2}\right] > 0.$$

Since $Q_L(0) < 0$ and $Q_L$ has exactly one positive root, we must have $0 < x_* < x_0$. It follows from Eq. (5) that

$$v_* = \frac{kB}{2} + x_* - x_*^2 > 0.$$

Therefore, $(x_*, v_*)$ is the unique physically accessible fixed point. □

**Lemma 5** (Local and global asymptotic stability)**.** For every $B, k > 0$, the unique physically accessible fixed point of the log-normal closure is locally asymptotically stable. Moreover, it is globally asymptotically stable in $\mathcal{D}_{\text{phys}}$ if and only if $b = kB(1 + 2B) \geq 1$.

*Proof.* We first establish local asymptotic stability. For a real $2 \times 2$ Jacobian matrix $J$, both eigenvalues have negative real parts if and only if $\operatorname{tr} J < 0$ and $\det J > 0$. At an equilibrium satisfying Eq. (5), direct differentiation and simplification give

$$\operatorname{tr} J_L = 10 - 12\frac{(x + kB/2)^2}{x^3} \tag{7}$$

and

$$\det J_L = 8 + 4kB + 12kB\frac{(x + kB/2)^2}{x^4} > 0. \tag{8}$$

Thus, it remains only to show that the trace is negative at $x_*$.

Let $x_{\text{tr}}$ denote the unique positive solution of

$$10x^3 = 12\left(x + \frac{kB}{2}\right)^2. \tag{9}$$

To see that this solution is unique, note that

$$\frac{\mathrm{d}}{\mathrm{d}x}\left[\frac{(x + kB/2)^2}{x^3}\right] = -\frac{(x + kB/2)(x + 3kB/2)}{x^4} < 0$$

for $x > 0$, while the expression in brackets decreases continuously from $+\infty$ to $0$. Hence the trace in Eq. (7) is negative precisely for $0 < x < x_{\text{tr}}$.

Using Eq. (9) to eliminate the cubic and quartic powers of $x_{\text{tr}}$ from $Q_L$ gives

$$\begin{aligned} Q_L(x_{\text{tr}}; k, B) = {} & \frac{48 + (384 + 120B)kB + 120(kB)^2}{25}\, x_{\text{tr}}^2 \\ & + \frac{48kB + (354 + 120B)(kB)^2 + 30(kB)^3}{25}\, x_{\text{tr}} \\ & + \frac{12(kB)^2 + (86 + 30B)(kB)^3}{25} > 0. \end{aligned} \tag{10}$$

Since $Q_L(0) < 0$ and $Q_L$ has exactly one positive root, Eq. (10) implies $x_* < x_{\text{tr}}$. Therefore $\operatorname{tr} J_L(x_*, v_*) < 0$, and Eq. (8) proves that the unique physically accessible fixed point is locally asymptotically stable.

We next turn to global stability. We first show that $b \geq 1$ is necessary. On the boundary $v = 0$, the second equation of Eq. (4) reduces to

$$\dot{v}\big|_{v=0} = b + 4x(x - 1) = b - 1 + 4\left(x - \frac{1}{2}\right)^2. \tag{11}$$

If $b < 1$, the right-hand side is negative for $x$ in an open interval containing $1/2$. By continuity of the vector field, trajectories starting sufficiently close to this boundary segment on the side $v > 0$ cross the boundary $v = 0$ and enter the negative-variance region. Hence $\mathcal{D}_{\mathrm{phys}}$ is not forward invariant, and global asymptotic stability in $\mathcal{D}_{\mathrm{phys}}$ is impossible.

It remains to prove sufficiency. Assume $b \geq 1$. Let $(x(\tau), v(\tau))$ be a trajectory starting in the interior of $\mathcal{D}_{\mathrm{phys}}$, and let $\tau_{\mathrm{e}}$ denote its first possible exit time from the interior. We first derive bounds for $0 \leq \tau < \tau_{\mathrm{e}}$. Since $v > 0$, the first equation of Eq. (4) gives $\dot{x} \leq kB + 2x - 2x^2$. Define

$$x_{\max} = \max\left\{x(0), \frac{1+\sqrt{1+2kB}}{2}\right\}.$$

The right-hand side is nonpositive at $x = x_{\max}$ and negative for $x > x_{\max}$. This clearly shows that $0 < x(\tau) \leq x_{\max}$ for $0 \leq \tau < \tau_{\mathrm{e}}$.

Using this bound in the second equation of Eq. (4), we obtain

$$\dot{v} \leq b + 4x_{\max}^3 + 4x_{\max}^2 + (4x_{\max} + 8)v - \frac{4v^3}{x_{\max}^3}.$$

The right-hand side tends to $-\infty$ as $v \to +\infty$. A standard comparison argument therefore gives a constant $v_{\max} > 0$ such that $0 < v(\tau) \leq v_{\max}$ for $0 \leq \tau < \tau_{\mathrm{e}}$. Thus, finite-time blow-up cannot occur before the trajectory reaches the boundary of $\mathcal{D}_{\mathrm{phys}}$.

We next show that the boundary $v = 0$ cannot be reached from the interior. If $b > 1$, Eq. (11) gives $\dot{v} > 0$ at every point on $v = 0$, and hence no trajectory starting with $v > 0$ can cross this boundary. The borderline case $b = 1$ requires slightly more care. In this case, Eq. (11) vanishes only at $P = (1/2, 0)$. At $P$, we have

$$\dot{x} = kB + \frac{1}{2} > 0, \quad \dot{v} = 0, \quad \partial_x \dot{v} = 0, \quad \partial_{xx} \dot{v} = 8.$$

Let the solution through $P$ satisfy $(x(\tau_0), v(\tau_0)) = P$. Along this solution, we have

$$\ddot{v} = \partial_x \dot{v}\, \dot{x} + \partial_v \dot{v}\, \dot{v},$$

and hence $\dot{v}(\tau_0) = \ddot{v}(\tau_0) = 0$. Differentiating once more and evaluating at $\tau_0$ gives

$$\dddot{v}(\tau_0) = 8\left(kB + \frac{1}{2}\right)^2 > 0.$$

Taylor expansion about $\tau_0$ therefore yields

$$v(\tau) = \frac{4}{3}\left(kB + \frac{1}{2}\right)^2 (\tau - \tau_0)^3 + o\big(|\tau - \tau_0|^3\big)\,. \tag{12}$$

Thus, the solution through $P$ lies in $v < 0$ for $\tau < \tau_0$ sufficiently close to $\tau_0$. If a trajectory starting in $v > 0$ were to reach $P$, uniqueness would force it to coincide locally with this trajectory, which is impossible. At every other point on $v = 0$, one has $\dot{v} > 0$. Therefore, the zero-variance boundary cannot be reached from the interior of $\mathcal{D}_{\mathrm{phys}}$ when $b \geq 1$.

It remains to exclude approach to the singular boundary $x = 0$. To this end, we choose $0 < \delta < \min\{x(0), 1\}$ and define $K = v_{\max}/\delta$. By decreasing $\delta$ if necessary, we may assume that $K \geq \max\{1, b/(kB)\}$. Suppose that the trajectory enters the strip $0 < x \leq \delta$ through the line $x = \delta$. Since $v \leq v_{\max} = K\delta$ at the entry point, the trajectory initially lies below the line $v = Kx$. On this line, direct substitution into Eq. (4) gives

$$\frac{\mathrm{d}}{\mathrm{d}\tau}(v - Kx) = b - kBK - 4K^3 + (-4 + 6K - 10K^2)x + (4 - 6K)x^2 < 0. \tag{13}$$

The inequality follows from $K \geq 1$ and $kBK \geq b$. Therefore, the vector field along $v = Kx$ points toward the region $v < Kx$, and the trajectory cannot cross this line from below. Hence $v(\tau) \leq Kx(\tau)$ whenever $0 < x(\tau) \leq \delta$.

Since $\delta < 1$, the first equation of Eq. (4) then gives $\dot{x} \geq kB - 2Kx$. Let

$$\varepsilon_x = \min\{\delta, kB/(4K)\} > 0.$$

Whenever $0 < x \leq \varepsilon_x$, we have $\dot{x} \geq kB/2 > 0$. Thus the trajectory cannot cross $x = \varepsilon_x$ from right to left and, in particular, cannot approach the singular boundary $x = 0$.

We can now conclude that the trajectory exists globally and remains in the physically meaningful region. Indeed, if $\tau_{\mathrm{e}} < \infty$, then the bounds obtained above imply that the trajectory remains in a compact subset of $\{(x, v) : x > 0\}$ whenever $0 \leq \tau < \tau_{\mathrm{e}}$. Since the vector field is continuously differentiable on this set, the solution can be extended beyond $\tau_{\mathrm{e}}$. On the other hand, the preceding arguments show that neither $v = 0$ nor $x = 0$ can be reached at $\tau_{\mathrm{e}}$, which is a contradiction. Hence $\tau_{\mathrm{e}} = \infty$. Moreover,

$$\varepsilon_x \leq x(\tau) \leq x_{\max}, \quad 0 < v(\tau) \leq v_{\max}, \quad \tau \geq 0.$$

Thus, every positive orbit is bounded and remains a positive distance away from the singular boundary $x = 0$.

It remains to establish convergence to the unique physically accessible fixed point. The interior of $\mathcal{D}_{\mathrm{phys}}$ is simply connected. Consider the Dulac function $\mathcal{B}(x, v) = x^{-5}$. Direct differentiation gives

$$\frac{\partial}{\partial x}(\mathcal{B}\dot{x}) + \frac{\partial}{\partial v}(\mathcal{B}\dot{v}) = x^{-6}\left[-5kB + 10(x^2 + v) - 12\frac{(x^2 + v)^2}{x^2}\right] < 0, \tag{14}$$

because $x^2 + v > x^2$ in the interior of $\mathcal{D}_{\mathrm{phys}}$. The Bendixson–Dulac criterion therefore excludes periodic orbits in the interior of $\mathcal{D}_{\mathrm{phys}}$ [4].

Every positive orbit is bounded and hence has a nonempty compact omega-limit set. This omega-limit set cannot intersect $v = 0$. Indeed, away from $P$, a boundary point has $\dot{v} > 0$, so its backward orbit immediately enters $v < 0$; when $b = 1$, the same conclusion at $P$ follows from Eq. (12). Hence the omega-limit set lies entirely in the interior of $\mathcal{D}_{\mathrm{phys}}$. If it contained no fixed point, the Poincaré–Bendixson theorem would imply that it is a periodic orbit [5], contradicting Eq. (14). Therefore, it must contain the unique physically accessible fixed point. Since this fixed point is locally asymptotically stable, it has an attracting neighborhood; membership in the omega-limit set forces the trajectory to enter this neighborhood, after which it converges to the fixed point.

Thus, when $b \geq 1$, every positive trajectory remains in $\mathcal{D}_{\mathrm{phys}}$ for all forward time and converges to the unique physically accessible fixed point. Combining the necessity and sufficiency arguments, we conclude that the unique physically accessible fixed point is globally asymptotically stable in $\mathcal{D}_{\mathrm{phys}}$ if and only if $b = kB(1 + 2B) \geq 1$. □

Lemmas 4 and 5 complete the proof of Theorem 2 in the main text.
